# The nitrogenase cofactor biogenesis enzyme NifB is essential for the viability of methanogens

**DOI:** 10.1101/2023.10.20.563283

**Authors:** Jasleen Saini, Ahmed Dhamad, Abaranjitha Muniyasamy, Andrew J. Alverson, Daniel J. Lessner

**Author notes:** Address correspondence to Daniel J. Lessner.

## Abstract

Dinitrogen (N_2_) is only bioavailable to select bacteria and archaea that possess the metalloenzyme nitrogenase, which reduces N_2_ to NH_3_ in a process called nitrogen fixation or diazotrophy. A long-term goal is to engineer diazotrophy into plants to decrease the use of nitrogen fertilizers, saving billions of dollars annually and greatly reducing nutrient pollution. This goal has not been realized, in part due to the inability to produce the nitrogenase metallocofactor within plants. Biogenesis of the cofactor requires NifB, a radical S-adenosy-L-methionine (SAM) enzyme that generates a precursor [8Fe-9S-C] cluster that matures into the final metallocofactor. Although maturation of nitrogenase is the only known function of NifB in bacteria, bioinformatic analyses reveal that NifB is conserved across methanogens, including those lacking nitrogenase, which suggests NifB functions outside of nitrogenase maturation. Indeed, several lines of evidence show that NifB is essential for viability of the model diazotroph, *Methanosarcina acetivorans*. First, CRISPRi repression was unable to abolish NifB production, whereas CRISPRi repression abolishes non-essential nitrogenase production. Second, unlike nitrogenase production, NifB production is not controlled by fixed nitrogen availability. Finally, *nifB* could not be deleted from *M. acetivorans* unless complemented *in trans* with *nifB* from other methanogens, including *Methanothrix thermoacetophila*, a species that lacks nitrogenase. Notably, *M. thermoacetophila* NifB supported diazotrophy in *M. acetivorans*, demonstrating that NifB from a non-diazotrophic methanogen produces the [8Fe-9S-C] cluster. Overall, these results link the metallocofactor biogenesis function of NifB to nitrogen fixation and methanogenesis, two processes of global importance.

**SIGNIFICANCE:** Methanogens directly impact life on Earth since they produce methane, a potent greenhouse gas, and are the principal archaea capable of nitrogen fixation, a process that requires nitrogenase. In this study, we demonstrate that NifB, an enzyme required to produce the metallocofactor in non-essential nitrogenase, is essential to the viability of methanogens. This identifies NifB as a new potential target in the goal of inhibiting methanogens to reduce methane emissions. The discovery that NifB functions outside of nitrogenase maturation will also aid efforts to engineer nitrogen fixation in plants, since NifB is a key factor to achieve this goal. Realization of these goals would have immense economic, environmental, and societal benefits.

## INTRODUCTION

Methanogenic archaea (methanogens) significantly impact life on Earth due to their ability to produce methane and to fix dinitrogen (N_2_) (1, 2). Methane is a critical link in the global carbon cycle but is also a potent greenhouse gas that significantly contributes to global climate change. Methane emissions from ruminant livestock are a major concern, estimated to accounting for an estimated 30% of all methane produced from anthropogenic sources (3). Methanogens and closely related methanotrophs are the only archaea capable of nitrogen fixation (diazotrophy), which is carried out by several lineages of bacteria. Archaeal and bacterial diazotrophs require molybdenum (Mo) nitrogenase to fix N_2_ into ammonia, a biologically reactive form of nitrogen (4, 5). Importantly, fixed nitrogen availability is the primary limiting factor in the growth of agricultural plants. Currently, the use of synthetic ammonia fertilizers is required to increase crop yields. Engineering diazotrophy in plants would eliminate our dependency on synthetic fertilizers, which would have enormous economic, environmental, and societal benefits (6-8).

Mo-nitrogenase is composed of two components: NifDK – the catalytic component that reduces N_2_, and NifH – the electron donor to NifDK (9-11). NifH contains a [4Fe-4S] cluster required for the ATP-dependent reduction of the [8Fe-7S] cluster (P-cluster) in NifDK. Electrons are then transferred from the P-cluster to the active site that contains a [7Fe-9S-Mo-C-homocitrate] cluster called the M-cluster or FeMo-co, which is the most complex metallocluster known in nature (12, 13). In addition to Mo-nitrogenase, some diazotrophs also contain vanadium (V) and iron-only (Fe) nitrogenases that are used when Mo is limiting. Each contains a cofactor similar to FeMo-co but with Mo replaced by V or Fe (i.e., FeV-co and FeFe-co) (14, 15). Despite our extensive understanding of the structure, function, and activity of nitrogenases, engineered plant diazotrophy has yet to be realized in part due to the inability to produce the complex metallocluster. Biogenesis of the cluster requires NifB, a radical S-adenosy-L-methionine (SAM) enzyme that generates an [8Fe-9S-C] cluster (NifB-co) that matures into FeMo-co, FeV-co, or FeFe-co (16-19). As such, understanding the functional properties of NifB is central to developing strategies to engineer diazotrophy in plants.

Nitrogenase maturation is the only known function of NifB in bacteria. In most bacteria, NifB contains a NifX domain that is thought to aid in the transfer of NifB-co to the maturase NifEN that serves as the scaffold for the synthesis of FeMo-co (4). However, all methanogen NifB proteins lack the NifX domain and, consequently, have proven more amenable to heterologous expression in a variety of hosts (20). Indeed, much of the understanding of the NifB structure and biochemistry, including the mechanism, has come from the characterization of recombinant methanogen NifB, with *Methanosarcina acetivorans* emerging as an important model system (17, 19-21). A recent breakthrough was the *in vitro* reconstitution of FeMo-co within Mo-nitrogenase using methanogen NifB expressed in the mitochondria of plant cells (22).

Although the functional properties of NifB from methanogens have been studied *in vitro* and within heterologous hosts, the *in vivo* function within methanogens is poorly understood. Recent work in *M. acetivorans* showed that all three nitrogenases support diazotrophy and that CRISPRi repression abrogates nitrogenase production, which in turn abolishes diazotrophy (23, 24). In contrast, CRISPRi repression of *nifB* was unable to abolish diazotrophy by *M. acetivorans* using Mo-nitrogenase (24). In this study, we present several lines of direct and indirect evidence, which show that NifB is essential for the viability of *M. acetivorans* and likely all methanogens. Thus, in addition to its function in nitrogenase maturation, NifB also serves an essential, albeit unknown, function in methanogens. This finding highlights important gaps in our understanding of a fundamental enzyme that holds the key to engineered nitrogen fixation in plants. This new insight can be leveraged to optimize efforts to engineer nitrogen fixation in plants and to also develop alternative strategies to mitigate methane emissions in livestock by targeting essential NifB in methanogens.

## RESULTS

### NifB is a universal feature of methanogen genomes

A total of 256 archaeal proteomes (245 methanogens, 11 non-methanogens) were assessed for the presence of NifB and nitrogenase with OrthoFinder. All methanogen proteomes contained at least one NifB protein that clustered into a single orthogroup (OG0000384), which included NifB from anaerobic archaea, including methanotroph and alkanotroph species (**Fig. 1 and Fig. S1**). Recently, the determined structure of NifB from *Methanobacterium thermoautotrophicum* revealed the amino acid ligands to the radical SAM (RS), K1, and K2 [4Fe-4S] clusters that are required to produce NifB-co. The RS cluster is coordinated by three cysteines, the K1 cluster by two cysteines and a histidine, and the K2 cluster by two cysteines and a histidine (21). Importantly, these residues are conserved across NifBs from all methanogen, methanotroph, and alkanotroph proteomes in our analysis, except for three species that have a cysteine in place of the K2 cluster histidine (**supplementary appendix**). A separate orthogroup (OG0000148) contained *M. acetivorans* NifD, VnfD, and AnfD, catalytic subunits from Mo-, V-, and Fe-nitrogenases, respectively. This orthogroup lacked nitrogenase-related subunits, including CfbD involved in cofactor F_430_ maturation in methanogens (25). Consequently, the presence of a protein in this orthogroup was used to infer the presence of nitrogenase within these proteomes. Despite the universal presence of NifB across methanogen proteomes, nitrogenase was present in 162 of them (∼67%) (**Fig. 1 and Fig. S1**). Thus, all methanogens possess a NifB that can likely produce NifB-co regardless of the presence or absence of nitrogenase. Since NifB in most bacteria has a C-terminal NifX domain (20), the prevalence of NifX in methanogens was also assessed. *M. acetivorans* contains three copies of NifX that fall within two separate orthogroups, designated here as NifX-1 and NifX-2. Analysis of the NifX-1 (OG0000173) and NifX-2 (OG0001145) orthogroups showed no correlation between NifX and the presence or absence of nitrogenase (**Fig. 1 and Fig. S1**).

**Figure 1.**
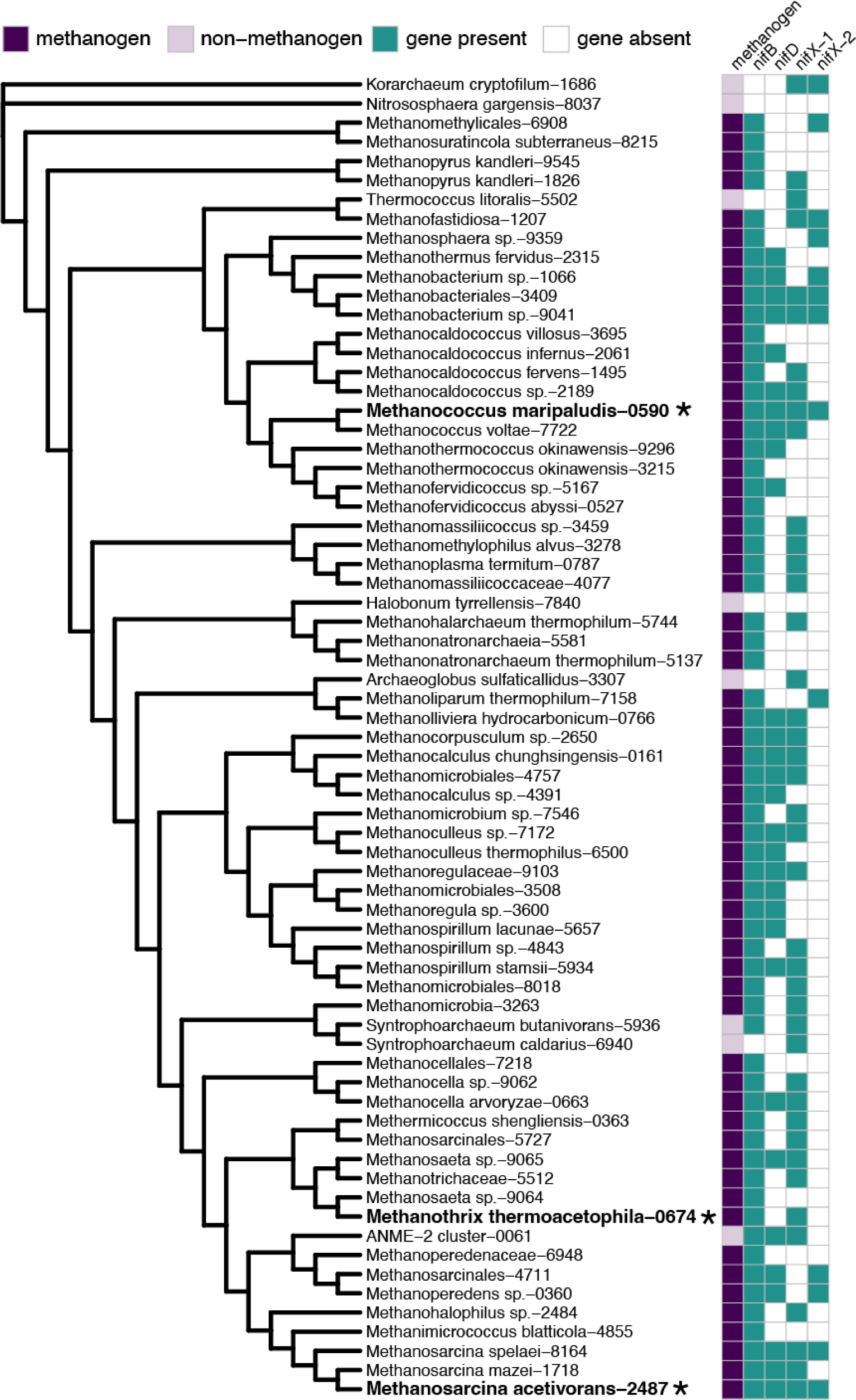
Phylogeny of methanogen and select non-methanogen archaea based on maximum likelihood analysis of a concatenated alignment of 104 low-copy orthologs. Species labels show the organism assignment from UniProt followed by the last 4 characters of the UniProt proteome identifier (Supplemental **Table S1**). For simplicity, genomes from the same taxonomic family and identical patterns of gene presence/absence were removed. The full tree is available in Supplementary **Fig. S1**.

CRISPRi repression of *nifB* with two gRNAs does not abolish diazotrophy by *M. acetivorans*.

Our previous results demonstrated that a single gRNA targeting dCas9 to the *M. acetivorans nif* operon (strain DJL74) abolishes diazotrophy, whereas the same approach with *nifB* (strain DJL105) only impaired Mo-dependent diazotrophy (24). This result was unexpected since both NifHDKEN encoded by the *nif* operon and NifB are essential for diazotrophy in bacteria (12, 13). To determine if the continued diazotrophy by strain DJL105 was due to inefficient CRISPRi repression, we generated two additional CRISPRi repression strains expressing a new gRNA (DJL140) or dual gRNAs (DJL141) (**Table 1**). The repression of *nifB* was analyzed in the new strains using qPCR to determine the relative abundance of *nifB* transcripts compared to DJL72, which lacks a guide RNA (**Fig. S2**). An approximate 90% reduction in *nifB* transcript abundance was observed in strain DJL141, whereas no reduction in transcript abundance was observed in strain DJL140. Repression of *nifB* in strain DJL141 is comparable to the repression previously observed in strain DJL105 (24).

**Table 1.**
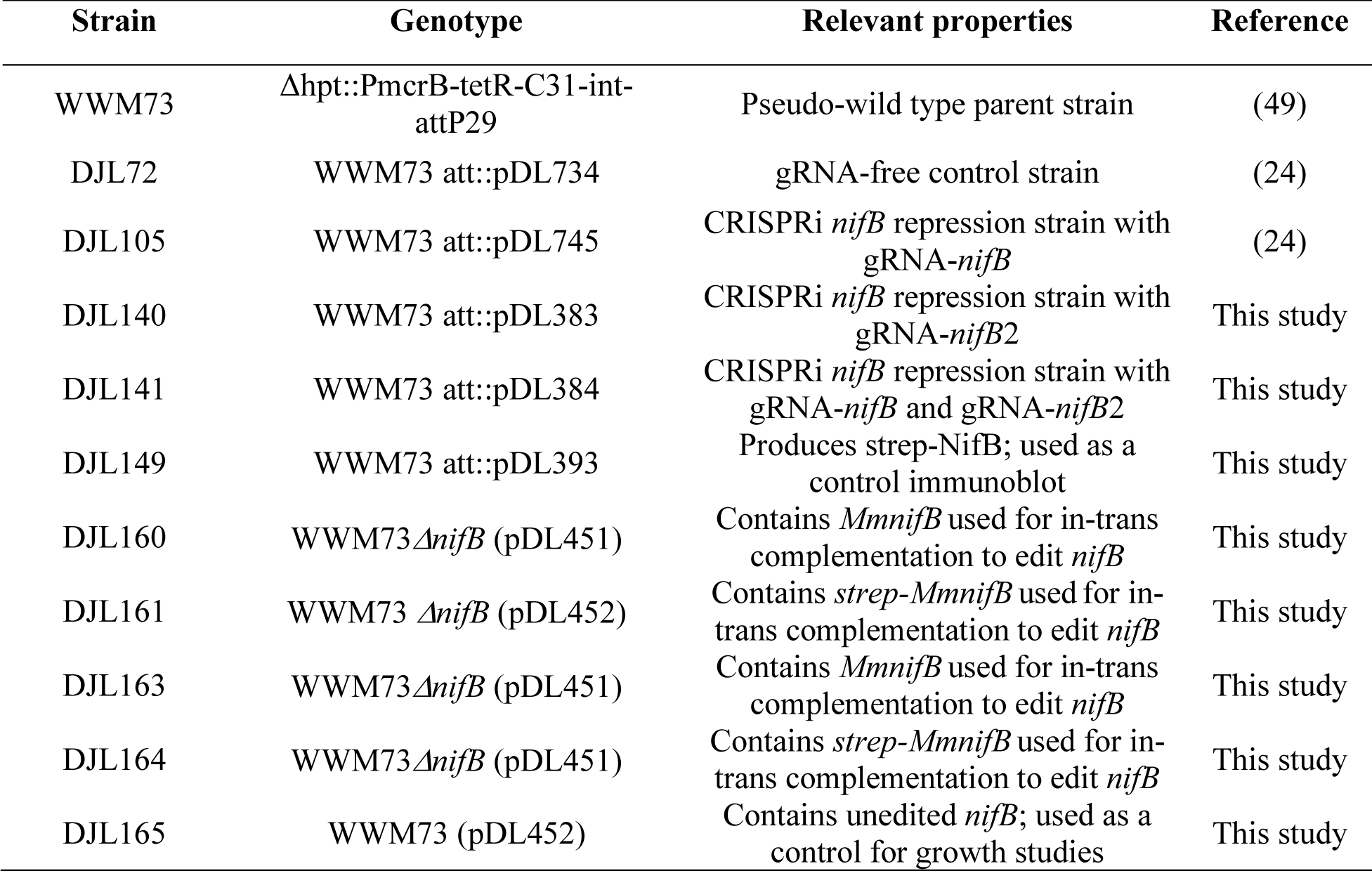
*M. acetivorans* strains used in this study.

| Strain | Genotype | Relevant properties | Reference |
| --- | --- | --- | --- |
| WWM73 | Δhpt::PmcrB-tetR-C31-int-attP29 | Pseudo-wild type parent strain | (49) |
| DJL72 | WWM73 att::pDL734 | gRNA-free control strain | (24) |
| DJL105 | WWM73 att::pDL745 | CRISPRi *nifB* repression strain with gRNA-*nifB* | (24) |
| DJL140 | WWM73 att::pDL383 | CRISPRi *nifB* repression strain with gRNA-*nifB2* | This study |
| DJL141 | WWM73 att::pDL384 | CRISPRi *nifB* repression strain with gRNA-*nifB* and gRNA-*nifB2* | This study |
| DJL149 | WWM73 att::pDL393 | Produces strep-NifB; used as a control immunoblot | This study |
| DJL160 | WWM73 Δ*nifB* (pDL451) | Contains *MmnifB* used for in-trans complementation to edit *nifB* | This study |
| DJL161 | WWM73 Δ*nifB* (pDL452) | Contains *strep-MmnifB* used for in-trans complementation to edit *nifB* | This study |
| DJL163 | WWM73 Δ*nifB* (pDL451) | Contains *MmnifB* used for in-trans complementation to edit *nifB* | This study |
| DJL164 | WWM73 Δ*nifB* (pDL451) | Contains *strep-MmnifB* used for in-trans complementation to edit *nifB* | This study |
| DJL165 | WWM73 (pDL452) | Contains unedited *nifB*; used as a control for growth studies | This study |

The growth of strains DJL140 and DJL141 with Mo was compared to strains DJL72 and DJL105 (**Fig. 2B**). All four strains grew similarly with NH_4_Cl (fixed nitrogen). Strain DJL140 also grew identically to strain DJL72 in medium lacking NH_4_Cl, consistent with the lack of *nifB* repression (**Fig. S2**). Without NH_4_Cl, strain DJL105 exhibits a slightly impaired growth phenotype as seen previously (24). However, growth of strain DJL141 without NH_4_Cl is more impaired but still not abolished. To determine whether NifB is still produced in strain DJL141, the level of NifB in lysates from cells of each strain was probed by Western blot (**Fig. 2C**). The intensity of the NifB band in strain DJL140 is similar to the control strain DJL72, whereas it is reduced in lysate from strains DJL105 and DJL141. These results indicate that continued Mo-dependent diazotrophy by strain DJL141 is likely due to reduced but not abolished NifB production. To further determine the effect of the reduced levels of NifB on Mo-dependent diazotrophy, the level of the Mo-nitrogenase catalytic subunit NifD was determined by Western blot (**Fig. S4**). No differences were detected, indicating impaired growth by strain DJL141 is not due to altered Mo-nitrogenase production but likely due to a limitation in FeMo-co biogenesis resulting from the reduced level of NifB.

**Figure 2.**
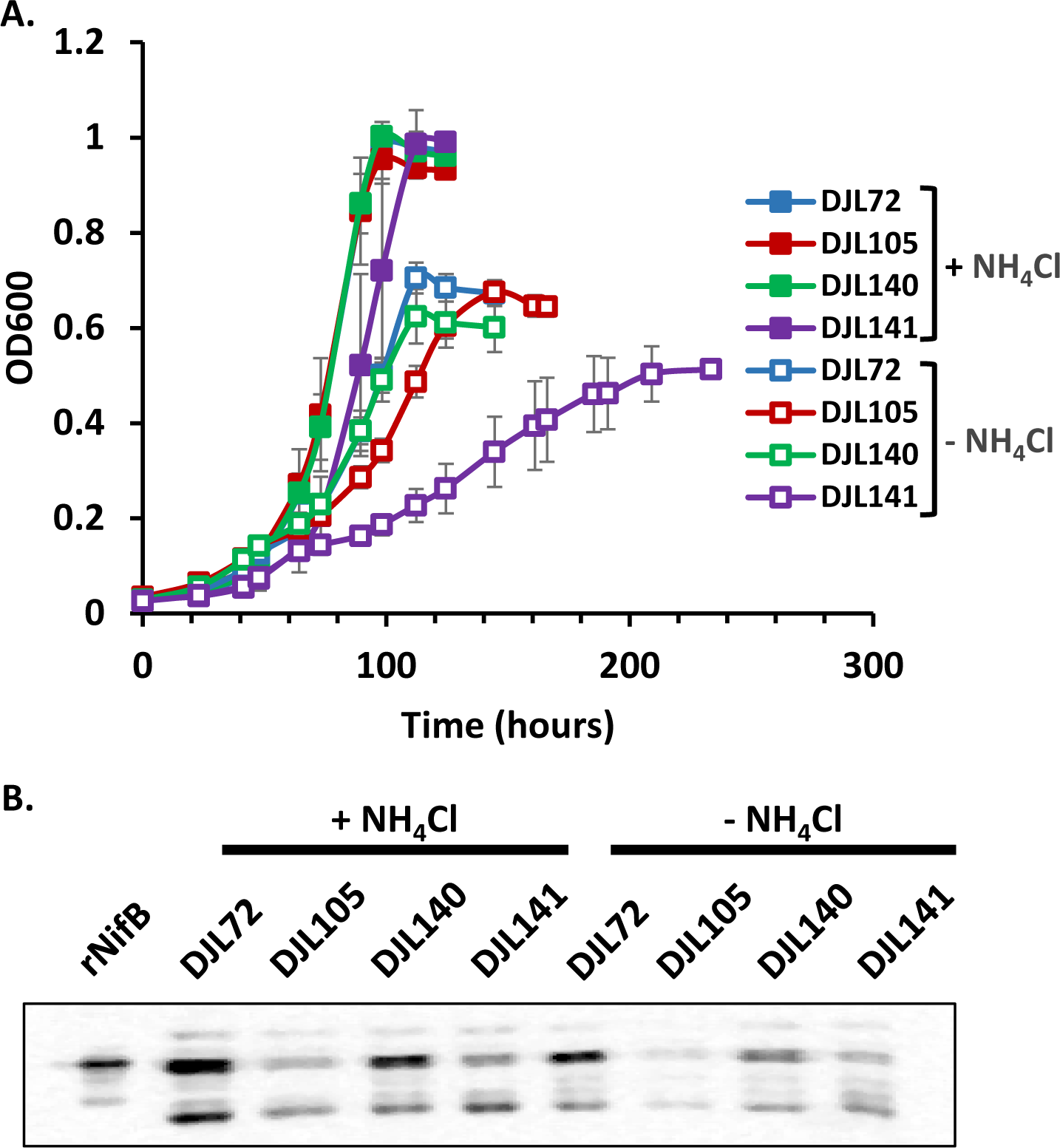
Growth and NifB expression analysis of *M. acetivorans* CRISPRi repression strains DJL105, DJL140, and DJL141 compared to control strain DJL72 (gRNA-free). (**A**) Comparison of growth in HS_DTT_ medium containing 125 mM methanol, 1 mM Na_2_S with or without NH_4_Cl. Error bars indicate mean ± SD for at least three biological replicates. (**B**) Western blot analysis of NifB protein abundance in cell lysates of *M. acetivorans* CRISPRi repression strains as in A. Lysate from *E. coli* expressing recombinant NifB (rNifB) was used as a specificity control. A protein loading control SDS-PAGE of lysates used for Western blot is shown in **Fig. S3**.

The impact of *nifB* repression during alternative nitrogenase usage was also tested by comparing the growth of strains DJL105, DJL140, and DJL141 to strain DJL72 in medium lacking Mo (Fe-only) or lacking Mo with added V (V + Fe). Unlike Mo-dependent diazotrophy, no significant difference was observed between the strains during Mo-independent diazotrophy (**Fig. S5**). To rule out loss of *nifB* repression during Mo-independent diazotrophy, the level of NifB was assessed by Western blot in cells growing in the Fe-only conditions (**Fig. S6**). Despite the lack of an impaired growth phenotype, the amount of NifB was still lower in strains DJL105 and DJL141 compared to control strain DJL72, similar to results with cells grown with Mo (**Fig. 2B**).

One notable observation of the Western blot in **Fig. 2B** is that NifB levels do not change in response to fixed nitrogen availability, unlike nitrogenases, which are only detected in cells grown with NH_4_Cl (24, 26). The relative transcript abundance of *nifB* in *M. acetivorans* parent strain WWM73 was determined by qPCR, and no difference in the transcript abundance is seen in cells grown with or without NH_4_Cl, consistent with observed protein levels (**Fig. S7**). These results show that NifB expression is not regulated at the transcriptional or translational level by the availability of fixed nitrogen, unlike the nitrogenases.

The CRISPR-Cas9 system is unable to delete *nifB* from the chromosome of *M. acetivorans*.

To determine whether NifB is essential for nitrogenase maturation, we used an established CRISPR-Cas9 system (27) to attempt to delete *nifB* from the chromosome of *M. acetivorans*. Our initial attempts used two gRNAs to generate two double-stranded DNA breaks necessary to delete the entire 972 bp gene. However, each deletion attempt (n = 3) resulted in the recovery of puromycin-resistant transformants that contained unedited *nifB*. To attempt to increase the efficiency of the editing of *nifB* by the CRISPR-Cas9 system, we next tried a single gRNA that would edit *nifB* such that the first ∼60% of the gene is deleted, including the region encoding the essential RS cluster (**Fig. S8**). A total of twelve puromycin-resistant transformants were screened for *nifB* editing by PCR using primers P15 and P16 that anneal outside of the upstream and downstream homology repair templates (**Fig. 3A**). Seven transformants produced a single PCR product consistent with edited *nifB*, four generated a single product consistent with unedited *nifB*, and one produced both products (**Fig. 3B)**. Sequencing of the PCR product from the putative edited *nifB* transformants confirmed the deletion of 60% of *nifB* as designed. However, PCR screening of the seven edited *nifB* transformants using primers P17 and P18 specific to *nifB* resulted in the production of a single PCR product consistent with the size of unedited (i.e., full-length) *nifB* (**Fig. 3C**), revealing the continued presence of unedited *nifB* (confirmed by sequencing the PCR product). Four additional transformations produced similar results, with all putative edited *nifB* transformants containing a mixture of unedited and edited *nifB* alleles based on PCR.

**Figure 3.**
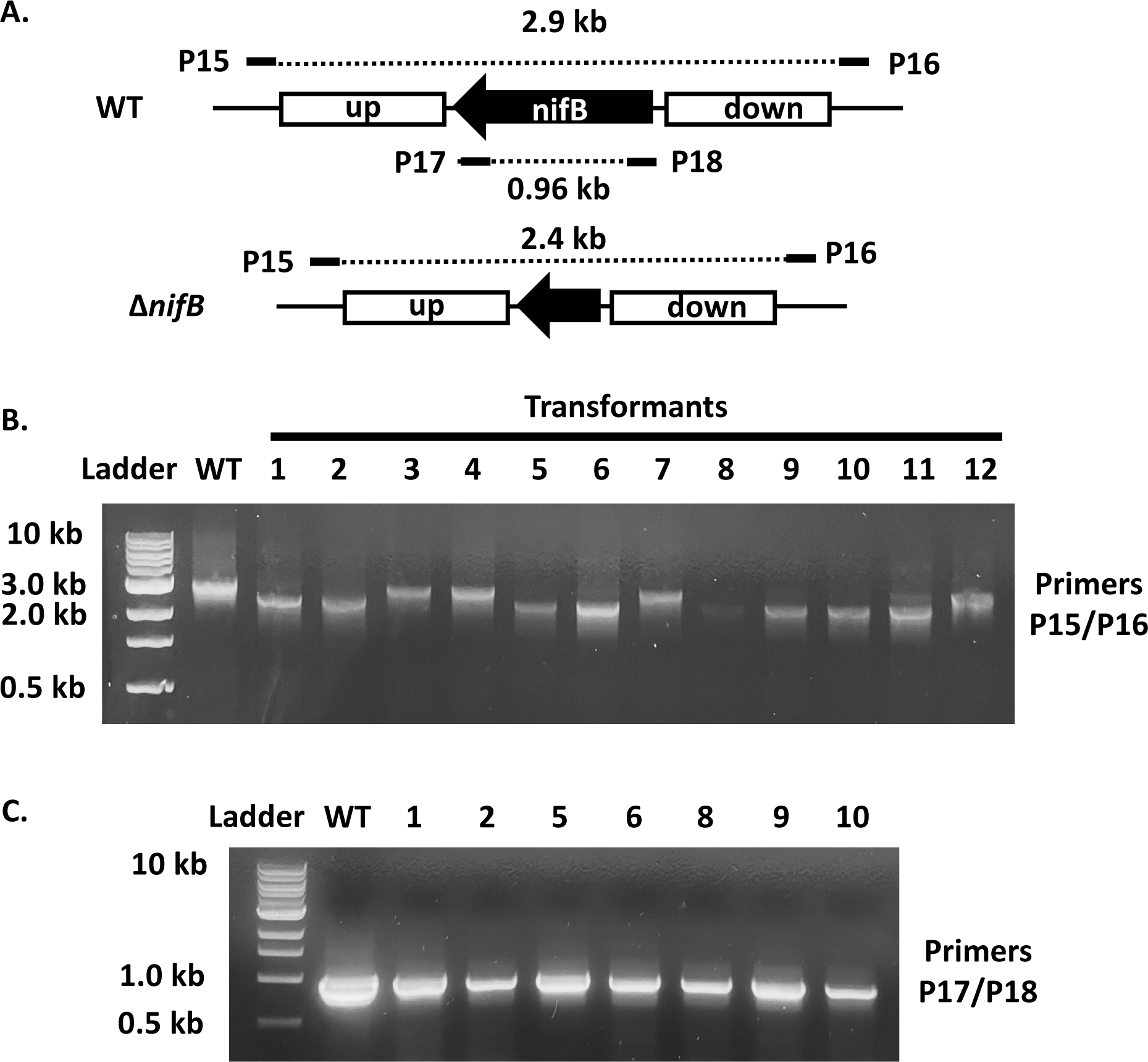
Screening of *M. acetivorans* transformants to assess CRISPR-Cas9 editing of *nifB*. (**A**) Schematic representation showing the size of predicted PCR products with indicated primers. (**B**) Gel image of PCR products with twelve transformants using primers P15 and P16 outside the upstream and downstream homology regions. (**C**) Gel image of PCR products with DNA from selected transformants using *nifB*-specific primers P17 and P18.

*M. acetivorans* is a known polyploid (28), leaving open the possibility that cells can simultaneously contain both unedited and edited *nifB* alleles. Moreover, gene conversion has been demonstrated in polyploid *Methanococcus maripaludis,* where an essential gene was targeted for deletion, but a resultant mixture of wild-type (WT) and mutant alleles, led to subsequent conversion to homozygous WT alleles (28, 29). A similar phenomenon could occur during our attempts to edit *nifB* in *M. acetivorans*. To test this, three WT-like transformants with only unedited *nifB* and three mutant-like transformants with only edited *nifB* as detected by PCR with primers P15 and P16 (**Fig. 4A**) were selected for gene conversion analysis. Each transformant was inoculated in standard medium with NH_4_Cl as the nitrogen source (a condition where NifB should not be necessary) and allowed to grow. Typical growth was observed for the WT-like transformants, but the mutant-like transformants exhibited a severe delay (>9 days) before the onset of growth and growth was more variable (**Fig. 4B**). Samples were taken at the end of growth and PCR was performed again with primers P15 and P16. Each mutant-like transformant sample now produced two bands, consistent with the appearance of unedited *nifB* alleles along with maintenance of edited *nifB* alleles. The transformants were passed in standard medium with NH_4_Cl a second time (**Fig. 4C**). Although the mutant-like transformants still exhibited impaired growth, it was much less severe than during the first passage. PCR was performed again, and only unedited *nifB* was detected in one mutant-like transformant, with the other two still containing a mixture of unedited and edited *nifB*. A third growth passage was performed, and the mutant-like transformants exhibited even less impaired and variable growth. PCR analysis of cells from the third passage resulted in the detection of only unedited *nifB* (**Fig. 4D**), demonstrating the conversion of edited *nifB* alleles to unedited *nifB* alleles. These results indicate that *nifB* is essential to the viability of *M. acetivorans*, such that despite the strong selective pressure of the CRISPR-Cas9 system to edit *nifB*, the cells retain unedited alleles which allows them to convert to a homozygous unedited *nifB* (WT) population.

**Figure 4.**
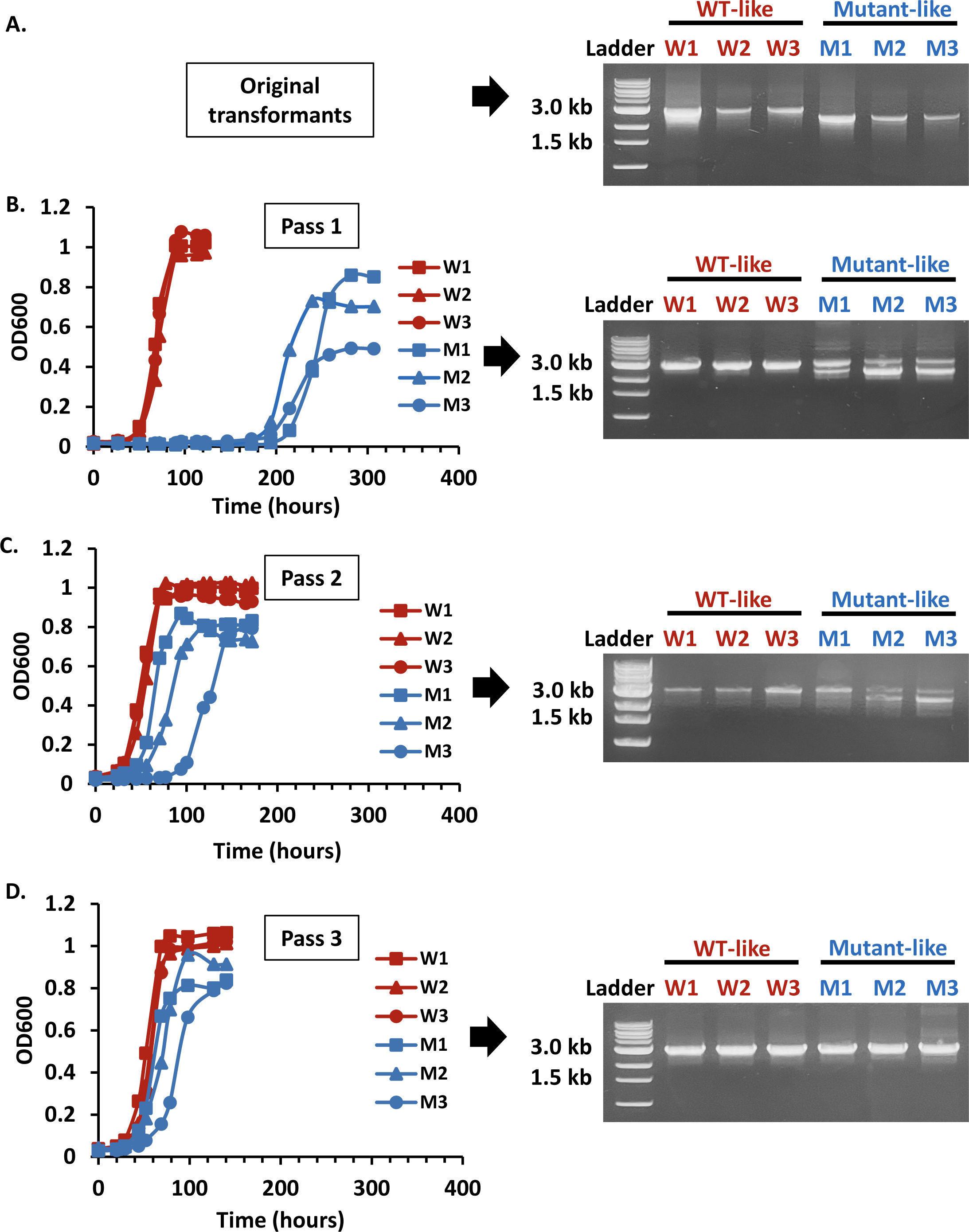
Comparison of the growth and the prevalence of edited *nifB* alleles of *M. acetivorans* wildtype (WT)-like and mutant-like *nifB* transformants during passage in HS medium containing NH_4_Cl. (**A**) PCR analysis using primers P15 and P16 of three WT-like and three mutant-like transformants as originally isolated. Comparison of the growth of the WT-like and mutant-like transformants and a subsequent repeat of PCR analysis after one (**B**), two (**C**), and three (**D**) passes in HS medium containing 125 mM methanol, 1 mM Na_2_S, 2 µg/ml puromycin, and 18 mM NH_4_Cl. Error bars indicate mean ± SD for three biological replicates.

How are the cells preventing the editing of *nifB* by the CRISPR-Cas9 system to allow gene conversion to occur? Each WT-like and mutant-like transformant harbors the CRISPR-Cas9 *nifB* editing plasmid, and all growth studies were performed in medium containing puromycin to maintain selection of the plasmid. Thus, loss of the plasmid cannot explain the ability of the cells to bypass Cas9-dependent editing of *nifB*. The most likely explanation is mutations arising within the gRNA, *cas9*, and/or *nifB* sequences that remove *nifB* as a target for Cas9-dependent editing. To test this hypothesis, the CRISPR-Cas9 *nifB* plasmid and PCR-amplified *nifB* were sequenced from the original transformant cells and from cells after the third passage. Notably, the original WT-like transformants contained no mutations within *cas9* and *nifB* but contained several mutations specifically within the gRNA sequence (**Fig. 5**), explaining why these cells never contained detectable edited *nifB*. The original mutant-like transformants that contained edited *nifB* (**Fig. 4**) had no mutations in *cas9*, *nifB*, or the gRNA revealing a functional CRISPR-Cas9 editing system and explaining the observed editing of the majority of *nifB* alleles. Sequencing of mutant-like cells after the third growth passage where all *nifB* alleles had converted to the unedited (WT) version still had no mutations in *cas9* or the gRNA (i.e., the CRISPR-Cas9 system is still functional). Instead, and remarkably, *nifB* now had a single point mutation in the third position of a glycine codon that is also the third nucleotide in the PAM sequence (**Fig. 5**), effectively removing *nifB* as a target for editing while maintaining the correct NifB amino acid sequence (i.e., still a codon for glycine). Therefore, the ability of the mutant-like cells to convert completely to unedited *nifB* (WT-like) is a result of strong selective pressure to retain functional NifB by removing *nifB* as a target of the gRNA for Cas9-dependent editing.

**Figure 5.**
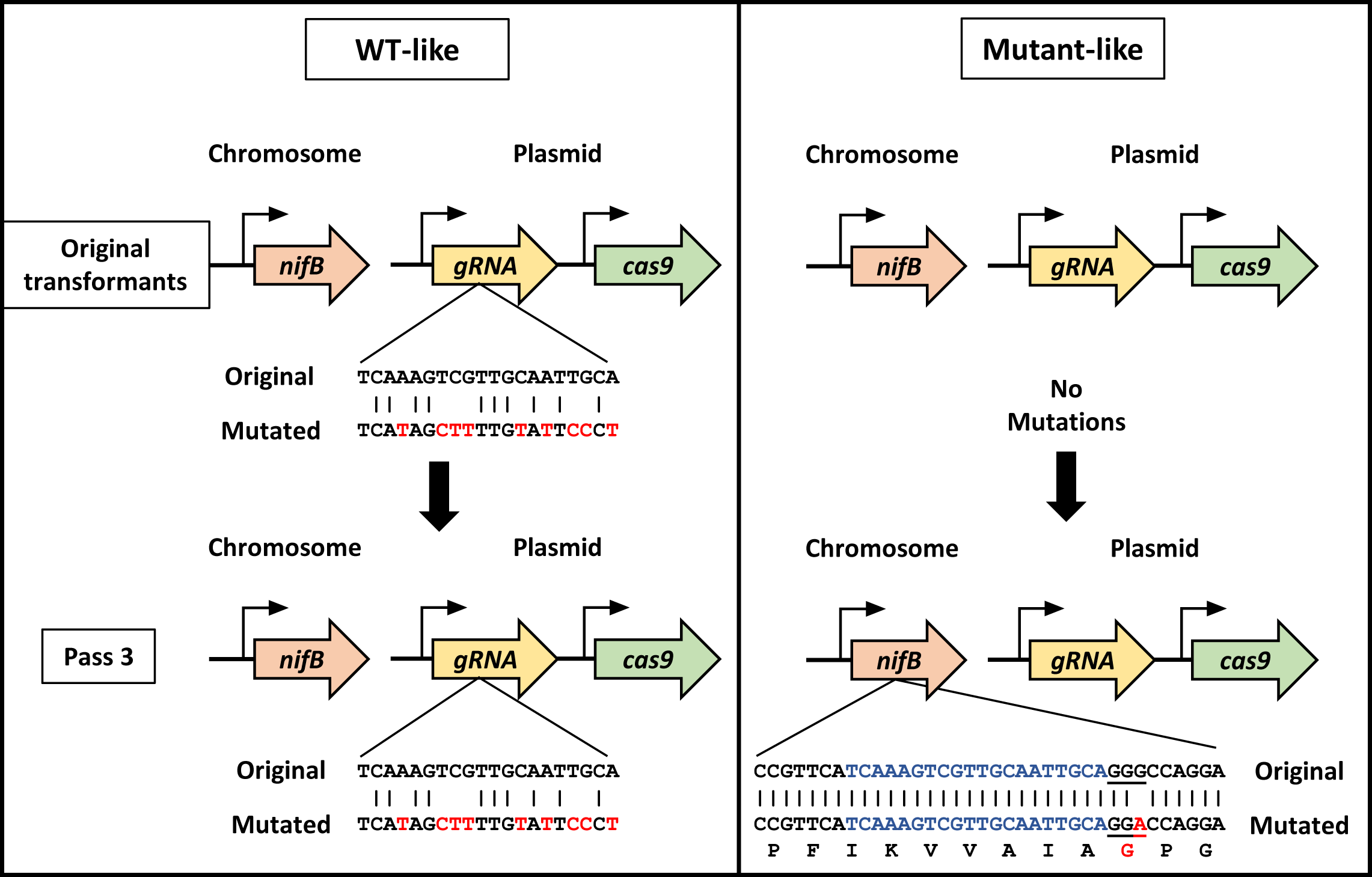
Model depicting the relevant mutations arising in the CRISPR-Cas9 system components (gRNA and *cas9*) or within *nifB* from WT-like and mutant-like transformants assessed in Fig. 4. Mutations in sequences are in red. The gRNA target sequence is in blue, and the PAM sequence is underlined.

### *M. acetivorans nifB* is edited in cells complemented with *nifB* from other methanogens

The inability to edit or delete *nifB* strongly suggests that NifB is essential for the viability of *M. acetivorans*. To further assess the essentiality of *nifB*, we attempted to delete *nifB* in cells complemented with *nifB* from the diazotrophic methanogen *Methanococcus mariplaudis* and the non-diazotrophic methanogen *Methanothrix thermoacetophila*. Specifically, *M. mariplaudis* NifB (MmNifB) and *M. thermoacetophila* NifB (MtNifB) with or without an N-terminal strep-tag were expressed at a different site within the CRISPR-Cas9 plasmid. *M. acetivorans* cells were transformed with CRISPR-Cas9 plasmids expressing either MmNifB or MtNifB and puromycin-resistant transformants for each were selected and analyzed as described above. Like the previous transformations, all the selected transformants still contained unedited *nifB*, even those with only edited *nifB* detected by PCR with primers P15 and P16. However, subsequent passage of the transformants containing only PCR-detectable edited *nifB* in medium with NH_4_Cl now resulted in the conversion to all edited *nifB* alleles. Importantly, unedited *nifB* is not detected by PCR, as shown for resultant strains DJL160 (MmNifB), DJL161 (strep-MmNifB), DJL163 (MtNifB) or DJL164 (strep-MtNifB), respectively (**Table 1 and Fig. S9**). Growth of the NifB-complementation strains was compared to strain DJL165, a puromycin-resistant WT-like strain harboring the CRISPR-Cas9 plasmid with MtNifB but is homozygous for unedited *nifB*. All four *nifB*-deletion strains exhibited growth in medium with NH_4_Cl similar to control strain DJL165 (**Fig. 6A**). Strains DJL160 and DJL161 expressing MmNifB and strep-MmNifB, respectively, also grew similarly to the control strain in medium lacking NH_4_Cl (**Fig. 6A**). However, strains DJL163 and DJL164 expressing MtNifB and strep-MtNifB, respectively, showed a severe delay prior to the onset of growth (**Fig. 6A**). Western blot analysis of lysates from cells of all four strains confirmed the absence of *M. acetivorans* NifB and the presence of strep-MmNifB in strain DJL161 (**Fig. 6B**). However, strep-MtNifB was not detected by Western blot in lysate from strain DJL164, despite the absence of *M. acetivorans* NifB (**Fig. 6B**). The delayed growth of strains DJL163 and DJl164 in the absence of NH_4_Cl is likely due to the low abundance of MtNifB in the strains since the Mo-nitrogenase catalytic subunit NifD abundance was similar to the control as determined by Western blot (**Fig. S11**). Nonetheless, these results demonstrate that both MmNifB and MtNifB can complement for the loss of NifB in *M. acetivorans* and are functional in nitrogenase maturation.

**Figure 6.**
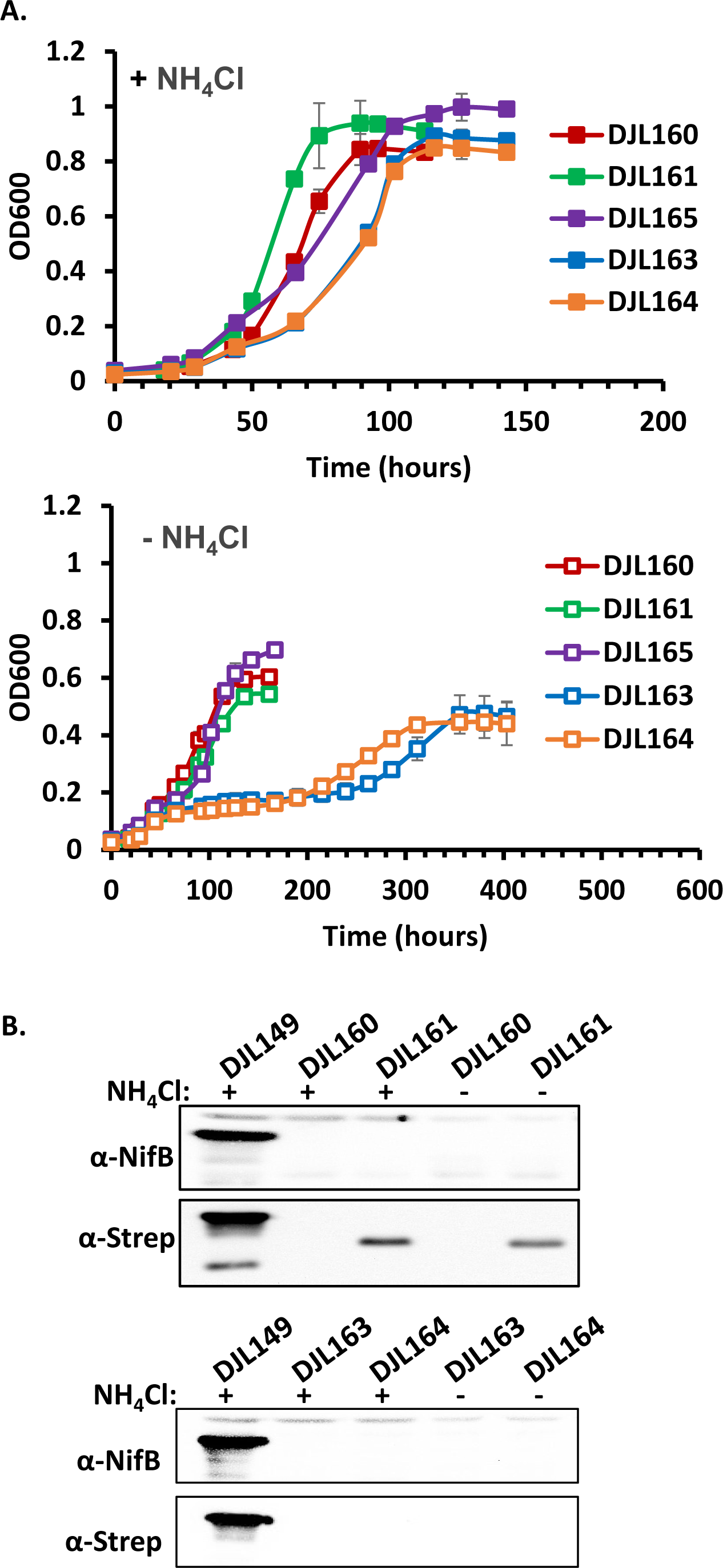
Growth and NifB expression analysis of *M. acetivorans* strains DJL160 (MmNifB), DJL161 (strep-MmNifB), DJL163 (MtNifB), and DJL164 (strep-MtNifB). (**A**) Comparison of the growth of each mutant strain to that of strain DJL165 (WT-like) in HS_DTT_ medium containing 125 mM methanol, 1 mM Na_2_S, and 2 µg/ml puromycin with or without NH_4_Cl. Error bars indicate mean ± SD for at least three biological replicates. (**B**) Western blot analysis using antibodies specific to NifB (α-NifB) or to strep-tag (α-Strep) of cell lysate from each strain as grown in panel A and compared to cell lysate from control strain DJL149 that expresses strep-NifB in addition to NifB (**Table 1**). Protein loading control SDS-PAGE of lysates used is shown in **Fig. S10**.

## DISCUSSION

Despite all current evidence indicating that nitrogenase maturation is the sole function of NifB, it has been previously suggested that NifB might serve a function outside of nitrogen fixation (20, 30). Our study clearly demonstrates that NifB does perform a separate function – one that is essential to the viability of *M. acetivorans* and likely all methanogens, given its universal presence. Methanogen NifB likely interacts with more proteins than previously known, a discovery that will facilitate efforts to optimize NifB-dependent biogenesis of the complex nitrogenase metallocofactor, one of the principal obstacles to engineering nitrogen fixation in plants. Further, our study identifies NifB as an essential factor in methanogens, which in the long term could be used as an antibiotic target to inhibit methanogen-derived methane emissions by livestock.

Why NifB is essential and what specific function it serves outside of nitrogenase maturation in methanogens is unclear. For nitrogenase maturation, NifB converts the K1 and K2 [4Fe-4S] clusters into a [8Fe-9S-C] cluster (NifB-co) by the RS cluster dependent insertion of carbide and the addition of a sulfur atom (17-19). Importantly, the RS, K1, and K2 cluster ligands are conserved in all methanogen NifB proteins, consistent with each capable of producing NifB-co, including those from methanogens lacking nitrogenase. Interestingly, the determined structure of MtNifB was missing the K2 cluster required for NifB-co production (17). However, MtNifB is capable of supporting diazotrophy in strains DJL163 and DJL164 that lack native NifB demonstrating that MtNifB produces NifB-co despite being in a methanogen that lacks nitrogenase. Given that MtNifB can produce NifB-co, it is likely that the essential function of NifB also involves the synthesis and transfer of NifB-co or a similar cluster. NifB is only known to transfer NifB-co to nitrogenase (AnfDGK) and maturase/scaffolds (NifEN and VnfEN) (13, 31), indicating another nitrogenase/maturase-related protein also requires NifB-co to carry out its essential function. Importantly, divergent members of the nitrogenase superfamily play critical roles in other processes, including within methanogens. For example, nitrogenase-like complexes participate in the biosynthesis of photosynthetic pigments (32, 33). Most notably, methanogens contain CfbCD, a nitrogenase-like system necessary for methanogenesis, although NifB is not known to be required. CfbCD functions in the biosynthesis of coenzyme F_430_, the tetrapyrrole cofactor of methyl-CoM reductase that catalyzes the methane-releasing step of methanogenesis (25). CfbC is homologous to the Fe-protein (NifH) of Mo-nitrogenase, and *in vitro* evidence supports CfbC as a homodimer with a single [4Fe-4S] cluster like NifH. Moreover, CfbC has been observed to form a weak complex with NifDK, the catalytic component of Mo-nitrogenase. Despite this interaction, CfbC + NifDK does not exhibit nitrogenase activity, indicating CfbC is unable to provide electrons to NifDK (34). CfbD is homologous to NifD but lacks a P-cluster or active site cofactor (e.g., FeMo-co). Instead CfbD is homomeric and contains a [4Fe-4S] cluster (35). Together CfbCD catalyzes the ATP-dependent six electron reduction of the isobacteriochlorin ring system of the intermediate Ni-sirohydrochlorin *a,c*-diamide (25). Although CfbCD is universally conserved in methanogens and is essential, it is unlikely that NifB functions directly with CfbCD since CfbD lacks an active site cofactor (i.e., there is likely no need for a [8Fe-9S-C] precursor cluster). Instead, CfbD directly binds the tetrapyrrole for reduction. Methanogens contain additional nitrogenase-like proteins, the function(s) of which are unknown (36, 37). A critical next step is to determine the essential function of NifB *M. acetivorans*.

Results from this study also provide new insights into methanogen genetics, particularly in relation to the use of CRISPR systems. The ability of transformants to switch from heterozygous (containing both edited and unedited *nifB* alleles) to homozygous (containing only unedited *nifB* alleles) is most likely a result of gene conversion (**Fig. 4**). In the absence of sexual reproduction, gene conversion can help offset the accumulation of deleterious mutations in asexual organisms (Muller’s Ratchet) (38). Gene conversion has been observed in other polyploid archaea, including *M. maripaludis* (55 genome copies) and *Haloferix volcanii* (20 genome copies) (28, 39, 40). *M. acetivorans* contains up to 17 genome copies when growing with methanol (28), the condition used in this study. Despite the efficiency of the *M. acetivorans* CRISPR-Cas9 system, it clearly cannot compete with the selective pressure to maintain unedited *nifB* because (i) rapid gene conversion due to spontaneous mutations quickly arising in the gRNA sequence in the CRISPR-Cas9 system likely accounts for only unedited *nifB* found in the original WT-like transformants (**Fig. 5**) and (ii) despite the mutant-like transformants having a functional CRISPR-Cas9 system that was able keep the number of copies of unedited *nifB* low initially, it only took a few generations for the transformants to convert from a heterozygous (edited and unedited *nifB*) to homozygous (unedited *nifB*) population (**Fig. 4**). Most remarkably, gene conversion in the mutant-like transformants is due to a single point mutation in the coding sequence of NifB that removed *nifB* as a target of Cas9 while maintaining functional NifB. These results reveal the plasticity and mutational buffering capacity of the polyploid genome of *M. acetivorans*.

Finally, the results provide new insight into the origins and evolution of nitrogenase, including the biogenesis of the complex metalloclusters. NifB likely originated in methanogens (4, 41), and recent evidence suggests nitrogenase likely evolved from ancestors that resemble maturases, the proteins that today function in nitrogenase cofactor assembly (e.g., NifEN) (42). These predecessor proteins evolved into modern enzymes harboring diverse metallocofactors and activities, such as nitrogenases and CfbCD, but there are likely others yet to be discovered. Indeed, our results indicate NifB serves as a broad metallocluster biogenesis protein in all methanogens. It not only functions in producing the precursor cluster to FeMo-co, FeV-co, and FeFe-co in nitrogenases, but it likely provides a precursor cluster to another nitrogenase-like enzyme that is essential to methanogens. In contrast, all current evidence indicates that NifB in bacteria is only involved in nitrogenase maturation. This could explain why most bacteria contain NifB with a NifX domain, whereas methanogen NifB proteins lack the NifX domain. NifX may provide more specificity to nitrogenase maturation in bacteria since NifX is thought to aid in transfer of NifB-co to NifEN (43). In methanogens, NifB likely needs to interact with nitrogenase-like proteins involved in the unknown essential function, in addition to nitrogenase and nitrogenase maturase proteins. Our results suggest that NifB likely originated in methanogens as an essential enzyme that was co-opted for nitrogen fixation, and subsequently transferred to bacteria as a nitrogenase-specific biogenesis enzyme (4). Overall, these results highlight important differences between nitrogen fixation in model bacteria and methanogens, and clearly emphasize that there is still much to learn about this process with the potential to help offset anthropogenic sources of nitrogen and carbon pollution.

## MATERIALS AND METHODS

### Comparative genomics

We downloaded 335 proteomes from the UniProt database, including methanogens and several representative non-methanogen relatives sampled based on the relationships in Auoad et al. (44). Proteomes were searched against the BUSCO Archaea database to evaluate completeness (45), and those with less than 90% complete or fragmented BUSCOs were removed, resulting in 256 proteomes (**Table S1**). Amino acid sequences were clustered into orthologous clusters (orthogroups) with OrthoFinder ver. 2.5.2 (46), and a species tree was inferred from 104 low-copy orthologs using the phylogenomic pipeline implemented in OrthoFinder. Presence of the diagnostic NifB Fe-S cluster ligands was evaluated with a Python script and validated by manual inspection of the multiple sequence alignment. The species tree and patterns of gene presence/absence were rendered with the R package ggtree (47). Data and scripts are available on Zenodo (10.5281/zenodo.10021994).

### *M. acetivorans* strains and growth

*M. acetivorans* strain WWM73 was the parent strain for all genetic experiments. All newly generated strains of *M. acetivorans* used in this study are listed in **Table 1**. *M. acetivorans* was grown in anoxic high salt (HS) medium containing 125 mM methanol and 1mM sulfide at 35°C as previously described (48). The medium was supplemented with 2 µg/ml puromycin when required. Growth experiments were performed in Balch tubes containing 10 ml of HS or Mo-deplete HS medium reduced with 1.5 mM DTT prepared as previously described (23). All strains were passed through Mo-deplete medium at least three times prior to the initiation of growth experiments. 125 mM methanol, 1 mM sulfide, 1 µM vanadate and 18 mM NH_4_Cl were added from anoxic sterile stock solutions prior to inoculation where indicated. Optical density was measured at 600 nm using a spectrophotometer.

### Construction of CRISPRi-repression strains

The *M. acetivorans* CRISPRi-dCas9 system was used as previously described (24) to generate strains DJL140 and DJL141 with *nifB* targeted for CRISPRi repression (**Table 1**). Detailed description of strain construction is provided in **supplementary note 1**.

### CRISPR-Cas9 editing of *nifB*

The *M. acetivorans* CRISPR-Cas9 system was used as previously described (27) with a few modifications to generate *M. acetivorans* strains DJL160-165 (**Table 1**). A detailed description of CRISPR-Cas9 editing of *nifB* in *M. acetivorans* and in-trans complementation with *nifB* from *M. maripaludis* and *M. thermoacetophila* NifB, can be found in **supplementary note 2**.

### Gene conversion analysis

Three WT-like transformants with unedited *nifB* and three mutant-like transformants with edited *nifB* based on PCR with primers P15 and P16 were selected for gene conversion analysis. Each transformant was grown in a Balch tube containing 10 ml HS medium supplemented with 18 mM NH_4_Cl, 3 mM cysteine, 125 mM methanol, 1mM sulfide and 2 µg/ml puromycin at 35 °C. Each transformant was passed in the same growth medium two more times using 200 µl of inoculum from the previous growth tube. Growth was monitored by measuring optical density at 600 nm using a spectrophotometer. Samples of cells (500 µl) were taken at the end of each growth passage and PCR was performed using primers P15 and P16.

### Gene expression analysis

*M. acetivorans* cells were harvested at mid-log phase (OD_600_ of 0.3-0.5) by anaerobic centrifugation. Pellets were resuspended in 1 ml Trizol reagent (Ambion, Life Technologies) and stored at -80°C. RNA was extracted using the Direct-zol RNA MiniPrep kit (Zymo Research) followed by DNase treatment using the DNA-free DNA Removal kit (Invitrogen, Thermo Fisher Scientific). cDNA was synthesized from 300 ng of RNA using the iScript Select cDNA Synthesis kit (Bio-Rad). Gene expression analysis was done by qPCR using cDNA (300-fold dilution) and SsoAdvanced Universal SYBR Green Supermix (Bio-Rad). Primers used for qPCR were designed using Geneious Prime software and purchased from IDT. Primers P22 and P23 were used for *nifB* expression analysis and primers P24 and P25 were specific to 16s rRNA (used as an internal control). The reactions were carried out in a CFX96 Real-Time PCR Detection system (Bio-Rad). The data were analyzed using ΔΔCq calculation method.

### Western blot analysis

Custom rabbit polyclonal antibodies specific for *M. acetivorans* NifB epitopes were purchased from GenScript. Mouse monoclonal antibody that recognizes the strep-tag epitope was purchased from Qiagen. Recombinant NifB expressed in *E. coli* was used as a positive control for Western blots. The detailed methods to construct the *E. coli* NifB expression strain are provided in **supplemental note 3**. For Western blot analyses, *E. coli* Rosetta DE3 cells containing pDL373 were grown in LB medium at 37 °C and induced with 50 mM IPTG at mid-log phase (OD_600_ of 0.5). The cells were harvested by centrifugation (8,000 x g for 5 minutes at 4 °C). The cell pellet was resuspended in buffer containing 50 mM Tris, 150 mM NaCl, 1 mM Benzamidine and sonicated 3 times on ice using a sonicator (QSonica) followed by aerobic centrifugation (8,000 x g for 5 minutes at 4 °C). The supernatant was removed, and the total protein concentration was determined using a Qubit protein assay kit (Molecular probes, Life Technologies). Each SDS-PAGE gel was loaded with 50 ng of supernatant protein as a positive control for western blots.

For western blot analyses using *M. acetivorans* lysate, cells were harvested at mid-log phase (OD_600_ of 0.3-0.5) by aerobic centrifugation (8,000 x g for 10 minutes at 4 °C). The cell pellet was resuspended in buffer containing 50 mM Tris, 150 mM NaCl, 1 mM Benzamidine and sonicated 3 times on ice using a sonicator (QSonica) followed by aerobic centrifugation (8,000 x g for 10 minutes at 4 °C). The cell lysate was collected, and the total protein concentration was determined using Qubit protein assay kit (Molecular probes, Life Technologies). Protein sample (10-15 µg) was resolved in a 12% SDS-PAGE gel and transferred to a PVDF transfer membrane (Thermo Scientific). The membrane was blocked for 15 minutes in buffer containing 50 mM Tris, 150 mM NaCl, 0.1% Tween (TBST), 5% milk and incubated overnight with the α-NifB, α-NifD, or α-strep antibody. The membrane was washed three times with goat anti-rabbit IgG antibody or goat anti-mouse IgG antibody (GenScript) for one hour followed by washing three times with TBST again. The membrane was developed with Clarity western ECL substrate (Bio-rad) and scanned using FlourChem 8900 imaging system (Alpha Innotech).

For western blot analyses of *M. acetivorans* in-trans complementation strains, control strain DJL149 expressing strep-NifB (**Table 1**) was used. The detailed methods to construct strain DJL149 are described in **supplementary note 4**.

## Acknowledgements

This work was supported in part by DOE Biosciences grant number DE-SC0019226 (DJL), NSF grant number MCB1817819 (DJL), and the Arkansas Biosciences Institute (DJL), the major research component of the Arkansas Tobacco Settlement Proceeds Act of 2000.

## Author contributions

JS, AD, and DJL conceived the study. JS and AD generated *M. acetivorans* mutant strains. JS performed growth studies, gene/protein expression studies, and gene conversion analyses. AM and AJA performed bioinformatic analyses. All authors contributed to data analysis and interpretation. JS prepared figures. JS and DJL wrote the manuscript. All authors reviewed the manuscript.

## Data availability

Data and scripts for computational analysis are available from Zenodo (10.5281/zenodo.10021994).

## SUPPLEMENTAL INFORMATION

### Supplemental methods

**Supplemental note 1: Further details on the construction of *nifB* CRISPRi-repression strains of *M. acetivorans.*** All primers and gBlocks were designed using Geneious Prime software and purchased from IDT. For the construction of CRISPRi-dCas9 plasmids pDL383 and pDL384, a gBlock containing gRNA-*nifB*2 (**Table S3**) was inserted into pDL734 and pDL745 (**Table S2**) at the HpaI site using a Gibson Assembly Ultra Master mix kit (Codex DNA) as per the manufacturer’s instructions. *E. coli* WM4489 competent cells (1) were transformed with the Gibson assembly reactions. Transformants were screened by PCR using primers P1 and P2 (**Table S4**) and the plasmid from a positive transformant for each was sequenced and named pDL383 (single *nifB* gRNA) and pDL384 (dual *nifB* gRNAs), respectively (**Table S2**). *M. acetivorans* strain WWM73 was separately transformed with pDL383 or pDL384 using liposome-mediated transformation as described (2). Transformants were selected on anaerobic HS agar plates containing 125 mM methanol and 2 µg/ml puromycin. Colonies were screened by PCR using two sets of primers P3/P4 and P5/P6 (**Table S4**) and a single positive colony for each was designated as strain DJL140 (pDL383) and strain DJL141 (pDL384) (**Table 1**). The strains were maintained in HS medium containing 125 mM methanol, 1 mM sulfide and 2 µg/ml puromycin.

**Supplemental note 2: further details on CRISPR-Cas9 editing of *nifB* in *M. acetivorans.*** gRNA-*nifB*\* (**Table S3**) was designed to guide Cas9 to cut within *nifB* such that the first ∼60% of the gene is deleted using homology dependent repair (**Fig. S8**). Primers, gRNA and gBlocks were designed using Geneious Prime and synthesized by IDT. The homology repair template downstream of *nifB* was amplified by PCR using primers P7 and P8 and the homology repair template upstream of *nifB* was amplified by PCR using primers P9 and P10 (**Table S4**). The homology repair templates and the gBlock containing gRNA-*nifB*\* were first assembled into BamHI-digested pUC18 using Gibson assembly and *E. coli* DH5α competent cells were transformed with the Gibson assembly reaction. Transformants were screened by PCR using primers P9 and P11, and a positive transformant was selected, the plasmid purified, and named pDL374 (**Table S2**). To move the homology repair template/gRNA-*nifB*\* fragment to the CRISPR-Cas9 plasmid pDL238, the fragment was amplified by PCR using primers P11 and P12 (**Table S4**) using Q5 polymerase and pDL374 as template. The PCR product was assembled into AscI-digested pDL238 using Gibson assembly, followed by transformation of *E. coli* WM4489 competent cells. Transformants were screened by PCR to confirm assembly and the resultant plasmid sequenced and named pDL377 (**Table S2**). pDL377 was retrofitted with pAMG40 using the Gateway BP Clonase II enzyme mix (Invitrogen) followed by transformation of *E. coli* WM4489 cells. Transformants were screened by PCR using primers P7 and P11, and the resultant plasmid was designated as pDL379. To attempt to edit *nifB*, *M. acetivorans* WWM73 cells were transformed with pDL379 using the liposome-mediated transformation method (2). Transformants were selected on anaerobic HS agar plates containing 125 mM methanol and 2 µg/ml puromycin. Colonies were transferred to 200 μl of HS medium containing 125 mM methanol, 1 mM sulfide and 2 µg/ml puromycin in a microtiter plate and grown in an anoxic jar at 35°C as previously described (3). An aliquot of cells (50 μl) were removed and screened for *nifB* editing by PCR using primers P15/P16 and P17/P18.

For in-trans complementation with *nifB* from *M. maripaludis* and *M. thermoacetophila* NifB, the *MmnifB* and *MtnifB* sequences (WP_011170602) were codon optimized using OPTIMIZER (http://genomes.urv.es/OPTIMIZER/) (4). MmNifB, strep-MmNifB, MtNifB, and strep-MtNifB were expressed from the CRISPR-Cas9 plasmid using the same promoter (*P_mcrB(tetO1)_*) and terminator used for Cas9 expression (5). gBlocks containing the promoter, codon optimized *MmnifB*, *strep-MmnifB*, *MtnifB* or *strep-MtnifB* and terminator fusion were designed using Geneious Prime software and synthesized from IDT. Plasmids pDL395-398 (**Table S2)** were constructed by digesting pDL377 with HpaI (NEB) and gBlocks were then introduced separately into digested pDL377 using Gibson Assembly Ultra Master mix kit (Codex DNA) as per the manufacturer’s instructions. *E. coli* WM4489 competent cells were transformed with the Gibson assembly reactions and transformants were screened by PCR using primers P1 and P2. Plasmids were purified and sequenced (Plasmidsaurus). Next, pDL395-398 were retrofitted with pAMG40 using Gateway BP Clonase II enzyme mix (Invitrogen) followed by transformation of WM4489 cells. The plasmids pDL395, pDL396, pDL397, and pDL398 retrofitted with pAMG40 were designated as pDL453, pDL454, pDL451, and pDL452, respectively. Transformants were selected and screened as described above. *M. acetivorans* strain WWM73 was then separately transformed with replicating plasmids pDL451-454 and transformants screened for *nifB* editing by PCR as described above. A transformant with complete editing of *nifB* was selected from each transformation and designated as strains DJL160 (pDL451), DJL161 (pDL452), DJL163 (pDL453), and DJL164 (pDL454). A WT-like transformant with unedited *nifB* due to mutations in the gRNA was designated as DJL165 and used as a control for the growth studies. The strains were maintained in HS medium containing 125 mM methanol, 1 mM sulfide and 2 µg/ml puromycin. Partial deletion of *nifB* was confirmed by sequencing PCR-amplified upstream and downstream regions around the deleted gene.

**Supplementary note 3: Construction of *E. coli* NifB expression strain as a positive control for Western blot analysis.** To construct the *E. coli* NifB expression strain, *nifB* was PCR amplified using *M. acetivorans* chromosomal DNA as the template and primers P17 and P18. PCR amplified *nifB* and pET28a were digested separately using NcoI-HF and BamHI-HF. Digested *nifB* and pET28a were ligated using T4 DNA ligase and *E. coli* DH5α competent cells were transformed with the ligation reaction. Transformants were screened by PCR using primers P17 and P18, and a positive transformant was selected, the plasmid purified, and named pDL373 (**Table S2**). Next, pDL373 was transformed into *E. coli* Rosetta DE3 competent cells.

**Supplementary note 4: Construction of *M. acetivorans* strain DJL149 expressing strep-NifB as a positive control for Western blot analysis of *M. acetivorans* in-trans complementation strains.** For the construction of strain DJL149, *nifB* was PCR amplified using *M. acetivorans* chromosomal DNA as the template and primers P20 and P21. PCR amplified *nifB* with strep tag at the C-terminus and pNB730 were digested separately using BamHI-HF and NdeI-HF. Digested *nifB* and pNB730 were ligated using T4 DNA ligase and *E. coli* DH5α competent cells were transformed with the ligation reaction. Transformants were screened by PCR using primers P20 and P21, and a positive transformant was selected, the plasmid purified, and named pDL393. *M. acetivorans* WWM73 cells were transformed with pDL393 using the liposome-mediated transformation method (2). Transformants were selected on anaerobic HS agar plates containing 125 mM methanol and 2 µg/ml puromycin. Colonies were screened by western blot for the presence of strep-NifB as described above and maintained in HS medium and designated as strain DJL149 (**Table 1**). The strains were maintained in HS medium containing 125 mM methanol, 1 mM sulfide and 2 µg/ml puromycin.

### Supplemental Tables

**Table S2.** Plasmids used in this study.

| Plasmid | Description | Reference |
| --- | --- | --- |
| pDN203 | *M. acetivorans* CRISPR-Cas9 system plasmid | (5) |
| pDL734 | pDN203 with Cas9 replaced by dCas9 | (6) |
| pDL745 | pDL734 with gRNA-*nifB* inserted at AscI site | (6) |
| pDL383 | pDL734 with gRNA-*nifB2* inserted at HpaI site | This study |
| pDL384 | pDL745 with gRNA-*nifB2* inserted at HpaI site | This study |
| pDL373 | pET28a with *strep-nifB* | This study |
| pDL238 | pDN203 without a gRNA | This study |
| pAMG40 | Retrofits with pJK027A-derived plasmids to allow replication in *M. acetivorans* | (7) |
| pNB730 | *M. acetivorans* expression plasmid | (8) |
| pDL374 | pUC18 with *nifB* -editing DNA (homology repair template + gRNA- *nifB* *) | This study |
| pDL377 | CRISPR-Cas9 plasmid (pDL238) with *nifB*-editing DNA | This study |
| pDL379 | pDL377 retrofitted with pAMG40 to allow replication in *M. acetivorans* | This study |
| pDL393 | pNB730 with *strep-nifB* | This study |
| pDL397 | pDL377 with *PmcrB(tetO1)-MmnifB* inserted at the HpaI site | This study |
| pDL398 | pDL377 with *PmcrB(tetO1)-strep-MmnifB* inserted at the HpaI site | This study |
| pDL451 | pDL397 retrofitted with pAMG40 to allow replication in *M. acetivorans* | This study |
| pDL452 | pDL398 retrofitted with pAMG40 to allow replication in *M. acetivorans* | This study |
| pDL395 | pDL377 with *PmcrB(tetO1)-MtnifB* inserted at the HpaI site | This study |
| pDL396 | pDL377 with *PmcrB(tetO1)-strep-MtnifB* inserted at the HpaI site | This study |
| pDL453 | pDL395 retrofitted with pAMG40 to allow replication in *M. acetivorans* | This study |
| pDL454 | pDL396 retrofitted with pAMG40 to allow replication in *M. acetivorans* | This study |

**Table S3.**
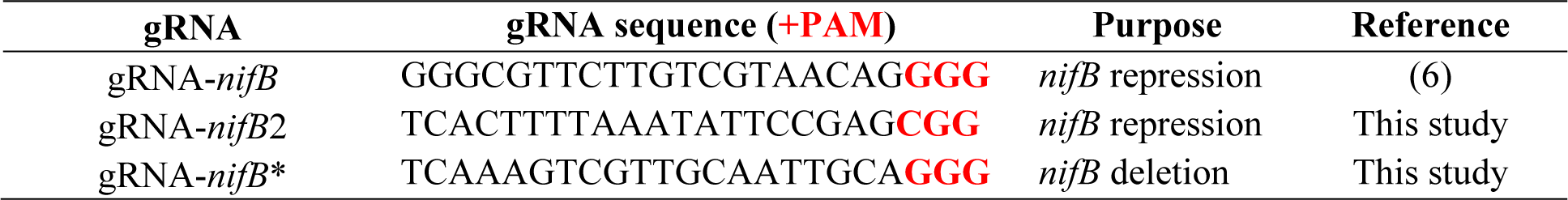
gRNAs used in this study.

**Table S4.**
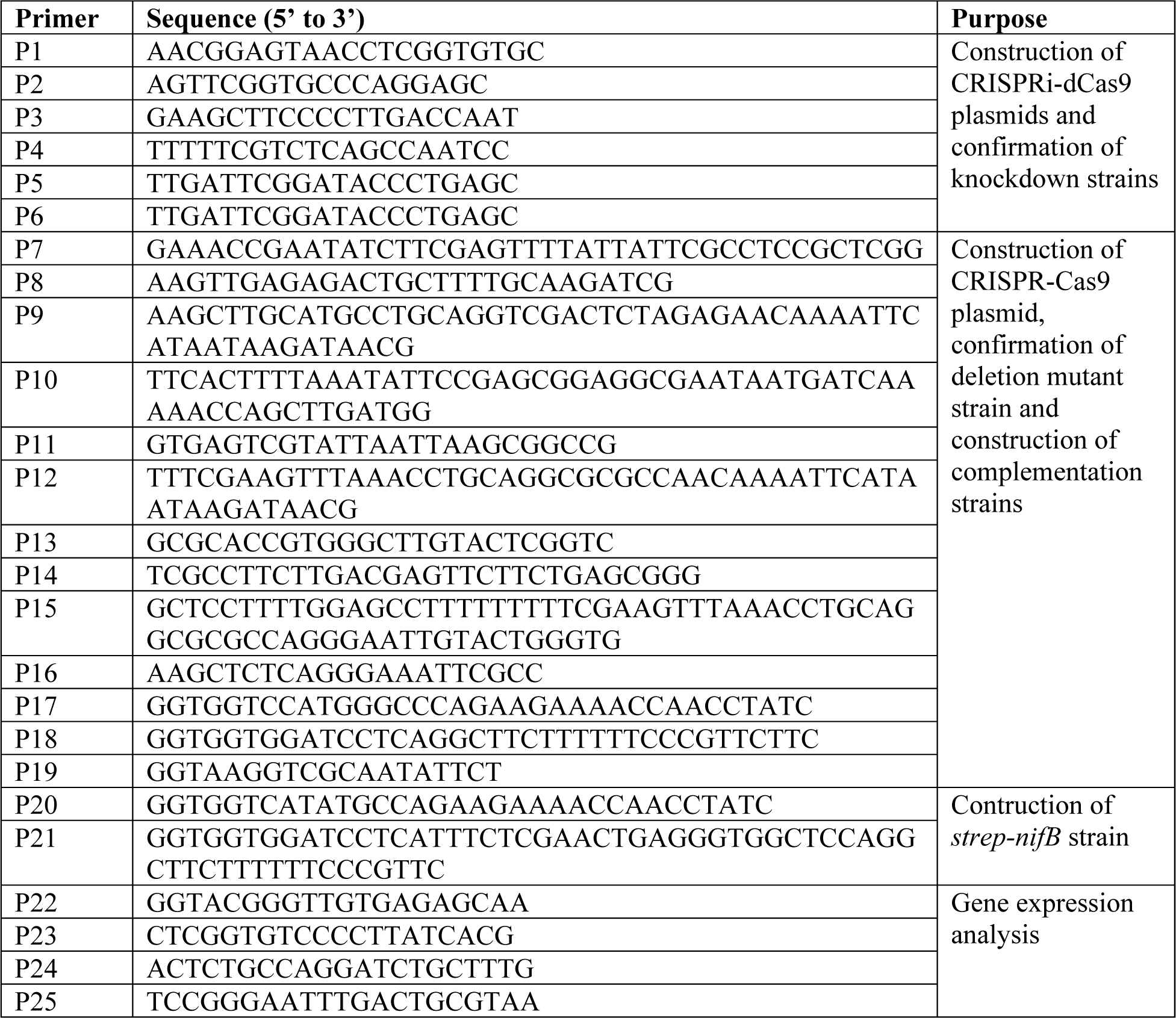
Primers used in this study.

---

**Figure S1.**
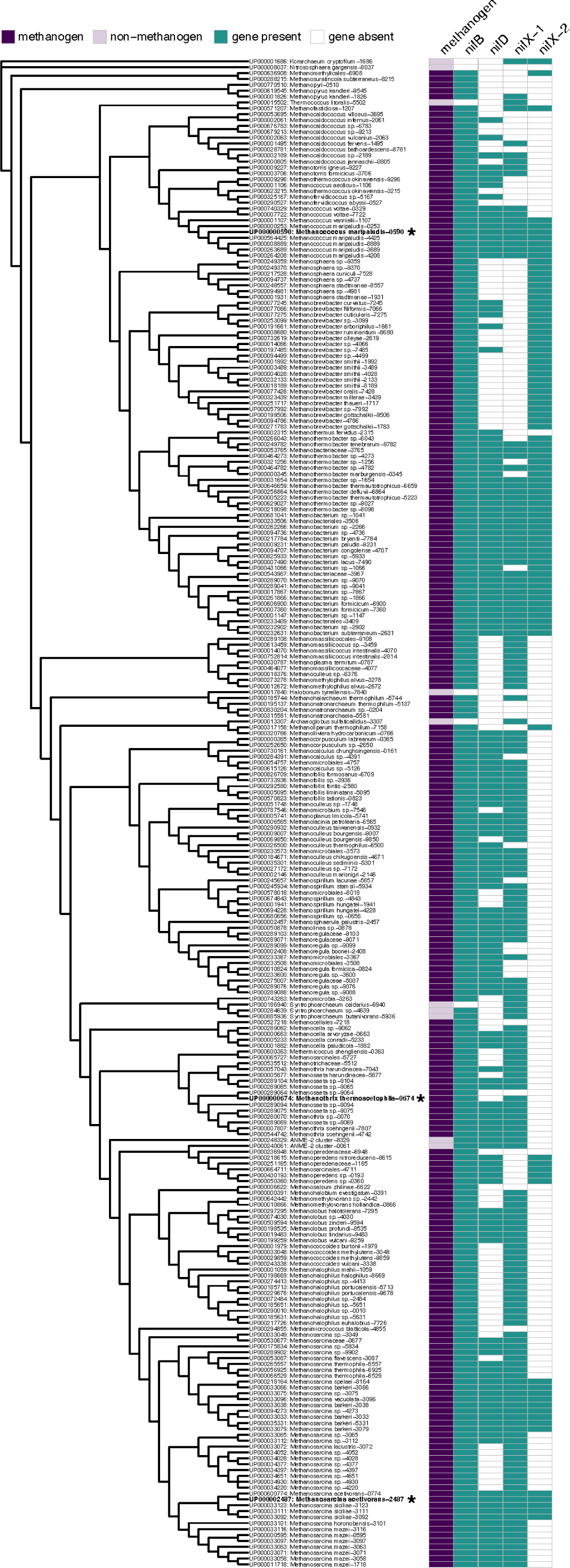
Phylogeny of methanogen and select non-methanogen Archaea based on maximum likelihood analysis of a concatenated alignment of 104 low-copy orthologs. Species labels show the UniProt proteome identifier, the organism assignment from UniProt, and the last 4 characters of the proteome identifier (Main text Fig. 1, Supplemental Table S1).

**Figure S2.**
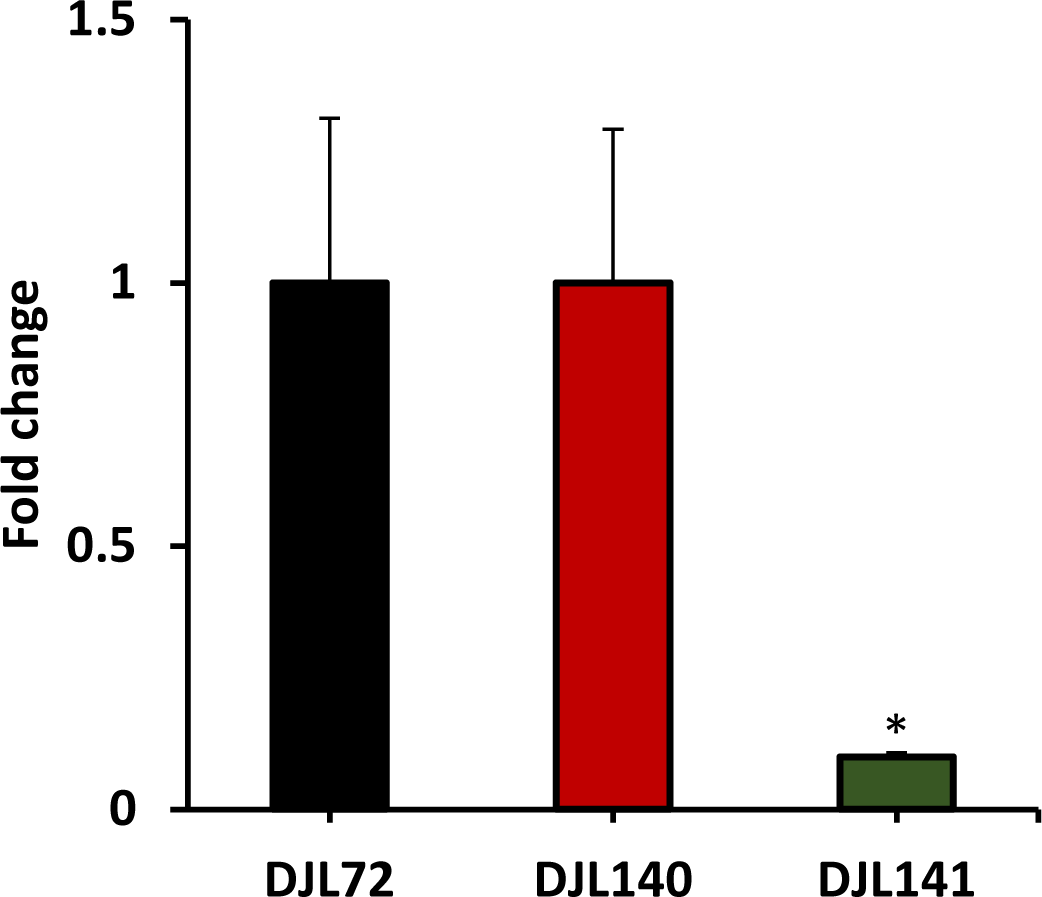
The relative transcript abundance of *nifB* in DJL140 and DJL141 compared to control strain DJL72 as determined by qPCR. Strains were grown in HS medium containing 125 mM methanol and 1 mM Na_2_S. Error bars indicate mean ± SD for three biological replicates. * p < 0.05

**Figure S3.**
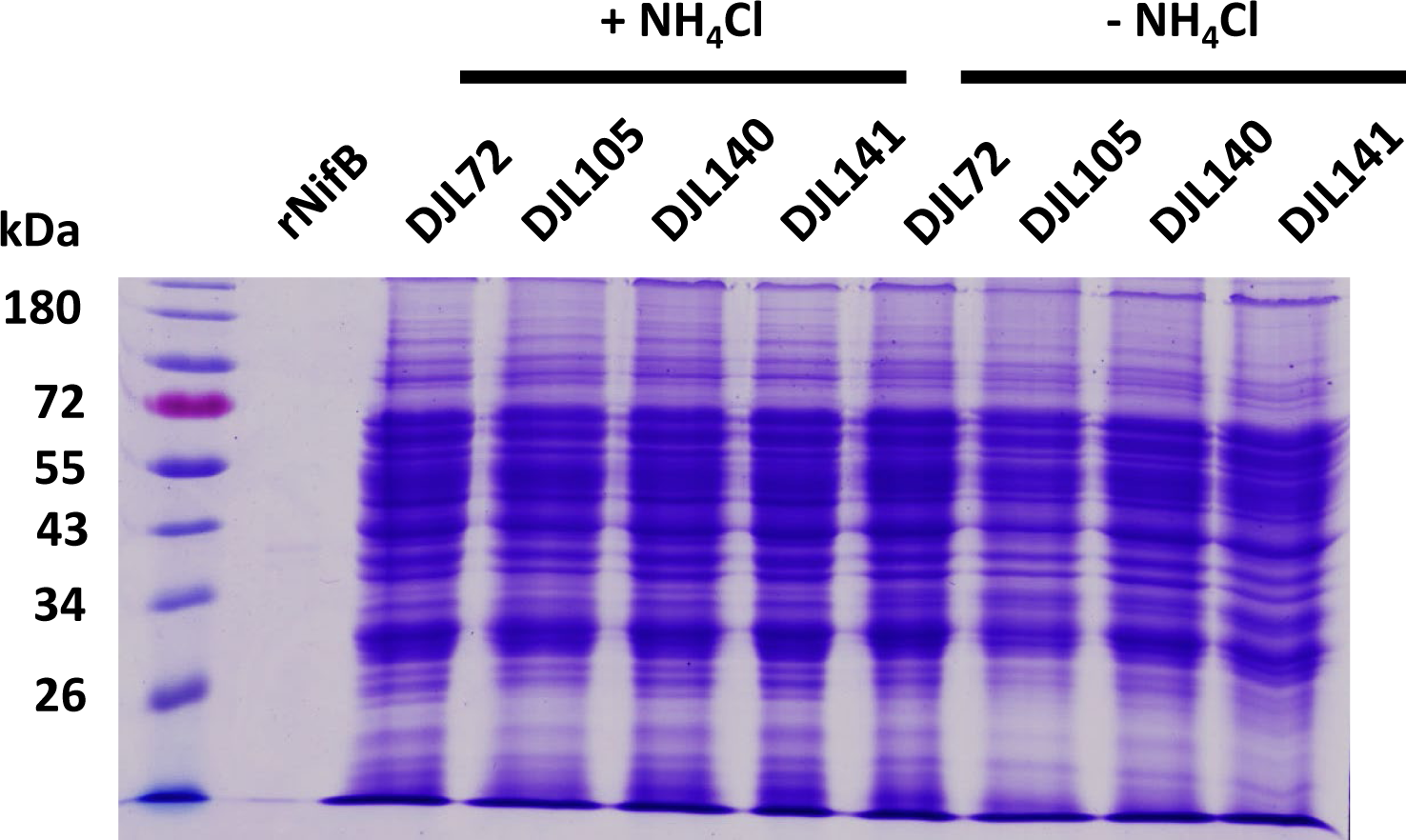
A protein loading control SDS-PAGE of lysates for the Western blot shown in Fig. 2.

**Figure S4.**
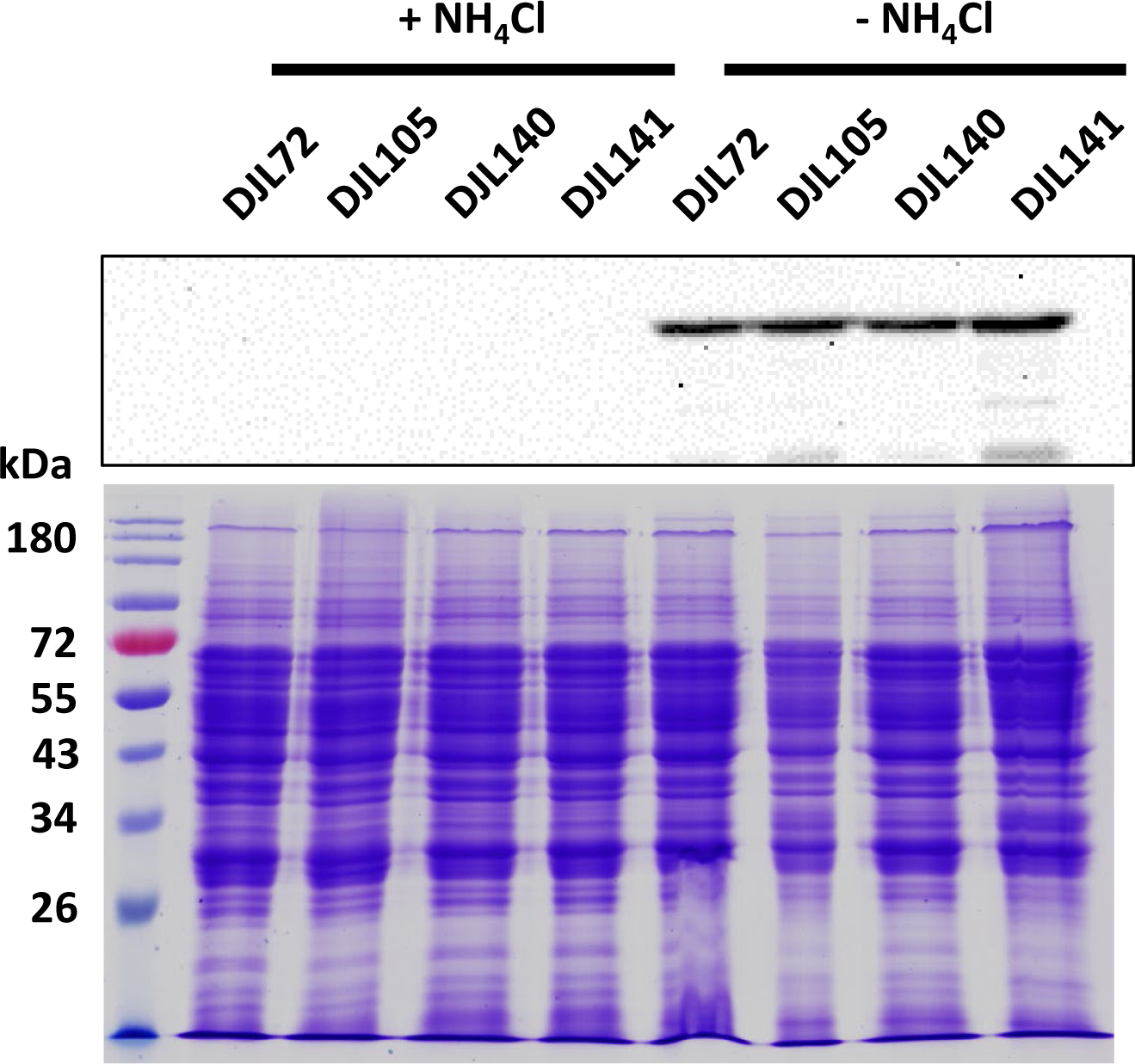
Western blot analysis of NifD abundance in cell lysates of *M. acetivorans* CRISPRi *nifB* repression strains. Strains were grown in HS_DTT_ medium containing 125 mM methanol and 1 mM Na_2_S with or without NH_4_Cl. Protein loading control SDS-PAGE of lysates used for is shown below Western blot.

**Figure S5.**
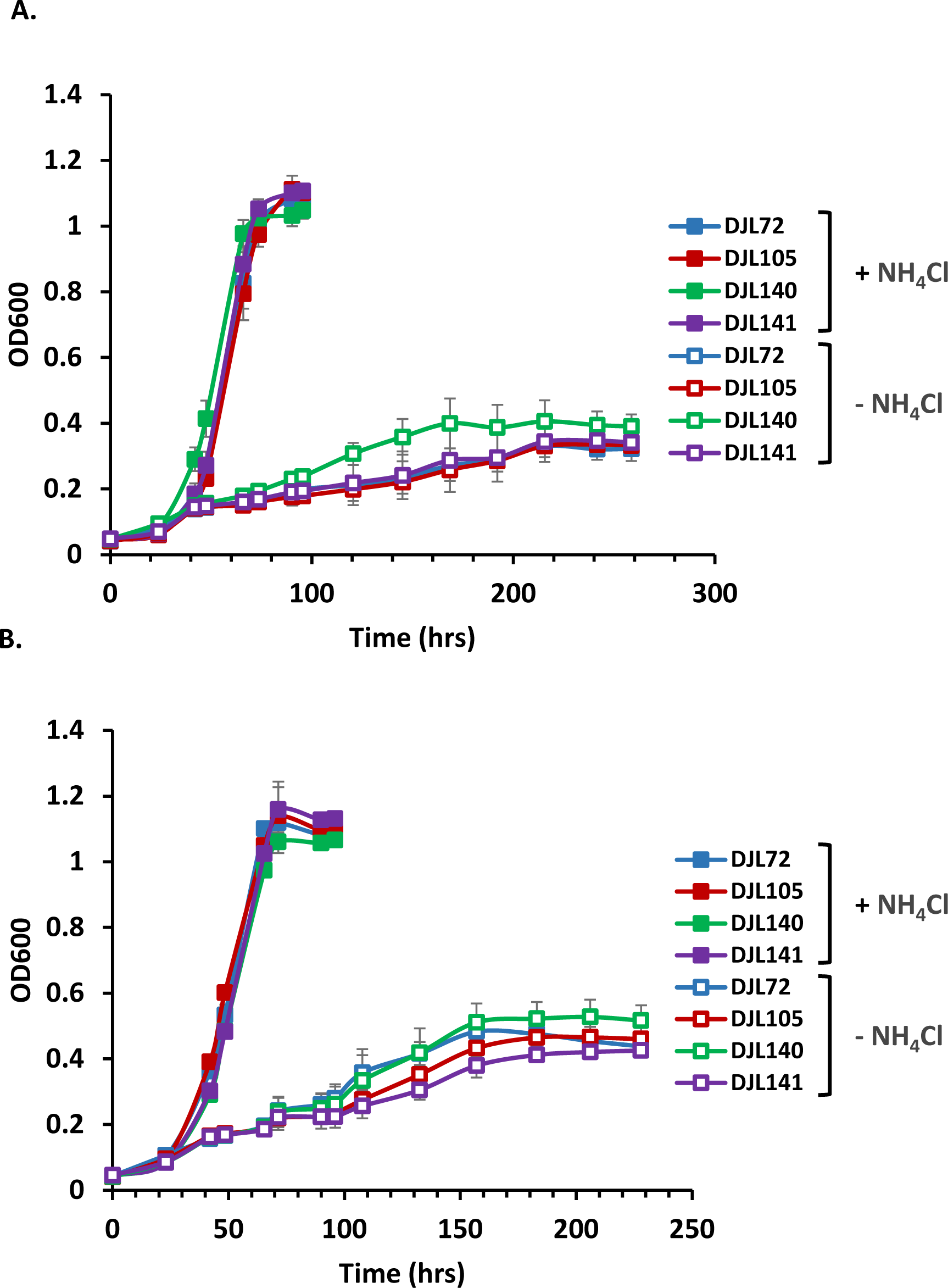
Comparison of the growth of *M. acetivorans* CRISPRi repression strains DJL105, DJL140, and DJL141 to the control strain DJL72 in HS_DTT_ medium depleted of Mo containing 125 mM methanol, 1 mM Na_2_S with or without NH_4_Cl. Medium was supplied (**A**) only Fe or (**B**) Fe+V. Error bars indicate mean ± SD for at least three biological replicates.

**Figure S6.**
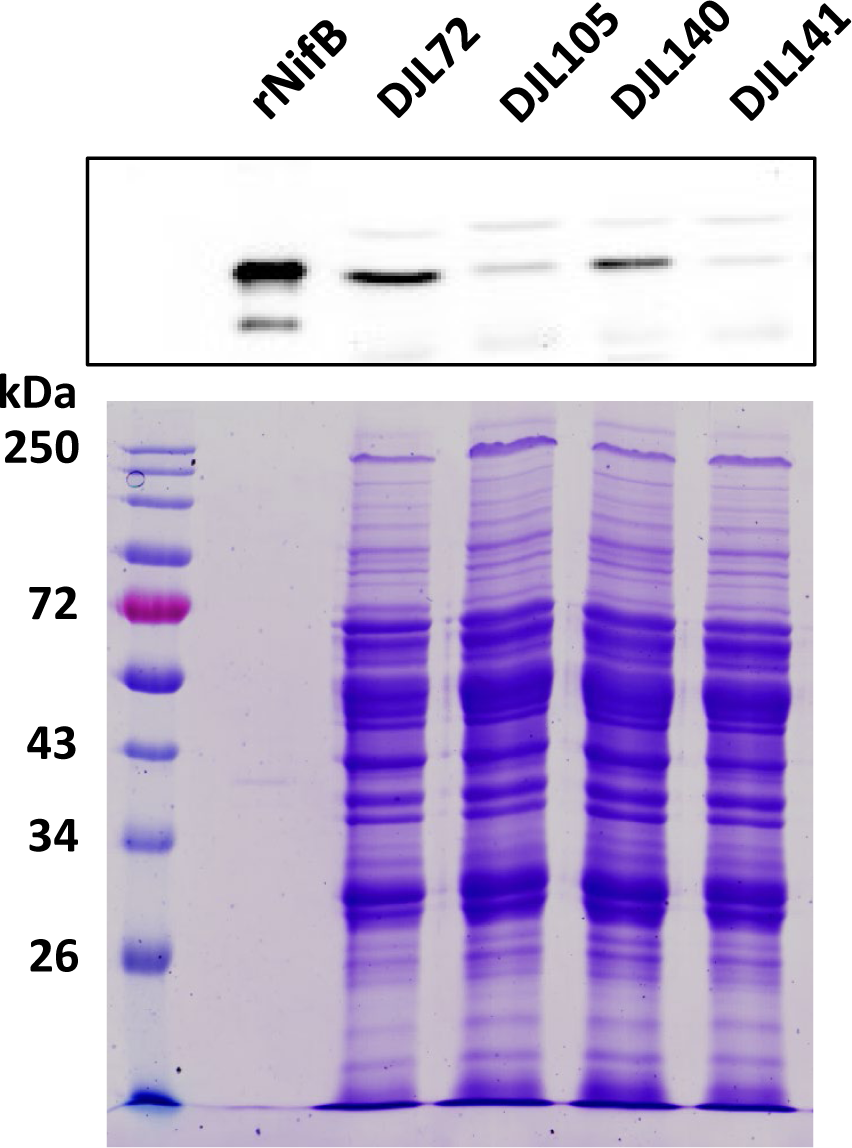
Western blot analysis of NifB abundance in cell lysates of *M. acetivorans* of CRISPRi *nifB* repression strains used for growth study in HS_DTT_ medium (Fe-only) containing 125 mM methanol and 1 mM Na_2_S without NH_4_Cl. (Top) Western blot analysis of lysates using antibodies specific to NifB. (Bottom) Protein loading control SDS-PAGE of lysates used for Western blot.

**Figure S7.**
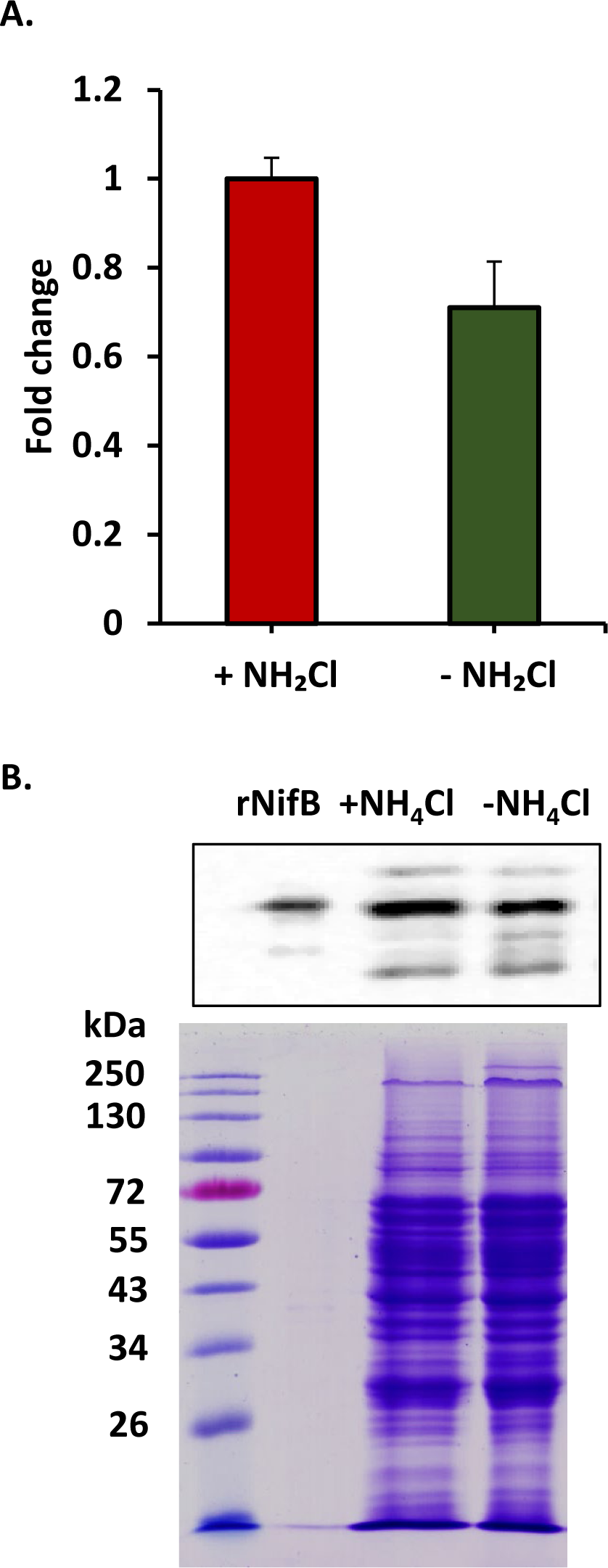
Effect of NH_4_Cl on expression of NifB in *M. acetivorans* strain WWM73(A) Relative transcript abundance of *nifB* in cells grown with and without NH_4_Cl as determined by qPCR. Strains were grown in HS_DTT_ medium containing 125 mM methanol and 1 mM Na_2_S. Error bars indicate mean ± SD for three biological replicates. * p < 0.05, ** p < 0.01, *** p < 0.001, **** p < 0.0001. (B, top) Western blot analysis of *M. acetivorans* cell lysates grown with or without NH_4_Cl using NifB-specific antibodies. (B, bottom) Protein loading control SDS-PAGE of lysates used for Western blot.

**Figure S8.**
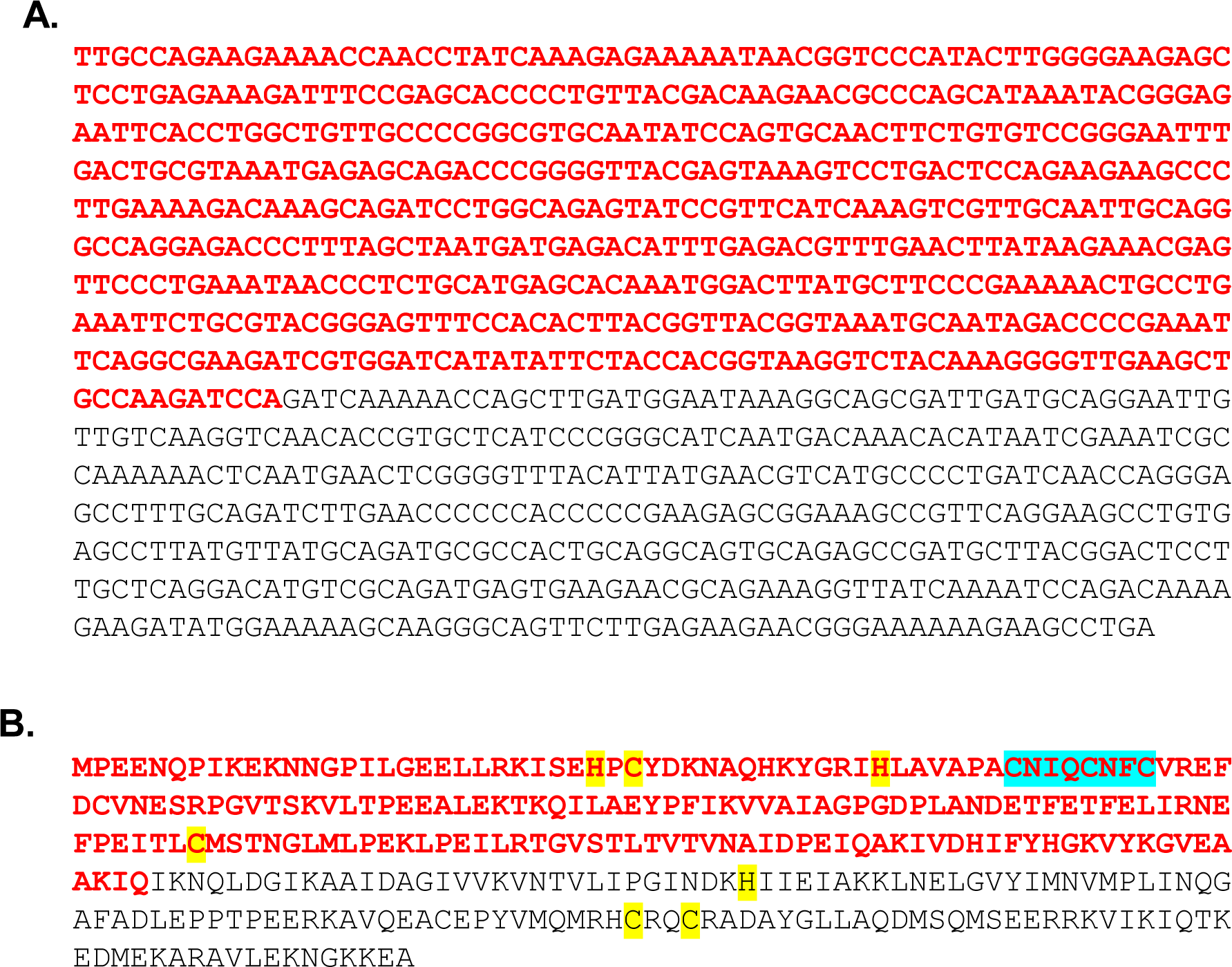
DNA (A) and protein (B) sequence of *M. acetivorans nifB*. Red denotes the designed deleted region using the CRISPR-Cas9 editing. In panel B, the SAM-cluster binding domain is highlighted in blue, and the conserved cysteine and histidine residues that bind the K1 and K2 [4Fe-4S] clusters are highlighted in yellow.

**Figure S9.**
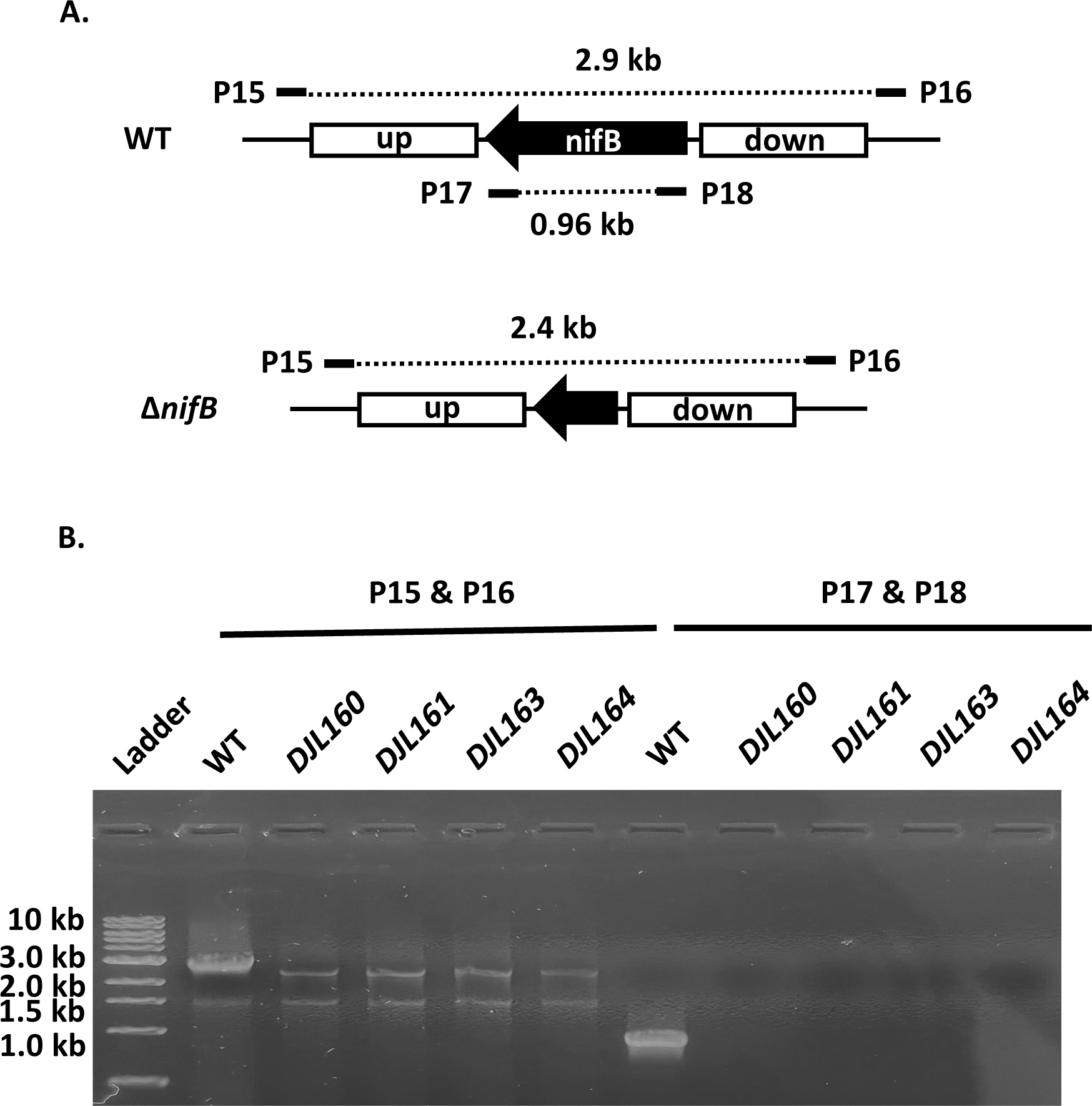
PCR analysis of *nifB* editing in *M acetivorans* strains DJL160, DJL161, DJL163, and DJL164 complemented with MmNifB, strep-MmNifB, MtNifB, and Strep-MtNifB, respectively, in comparison to WT using three different primer sets. (**A**) Schematic representation showing the size of predicted PCR products with the indicated primers. (**B**) Gel image of PCR products with indicated primers.

**Figure S10.**
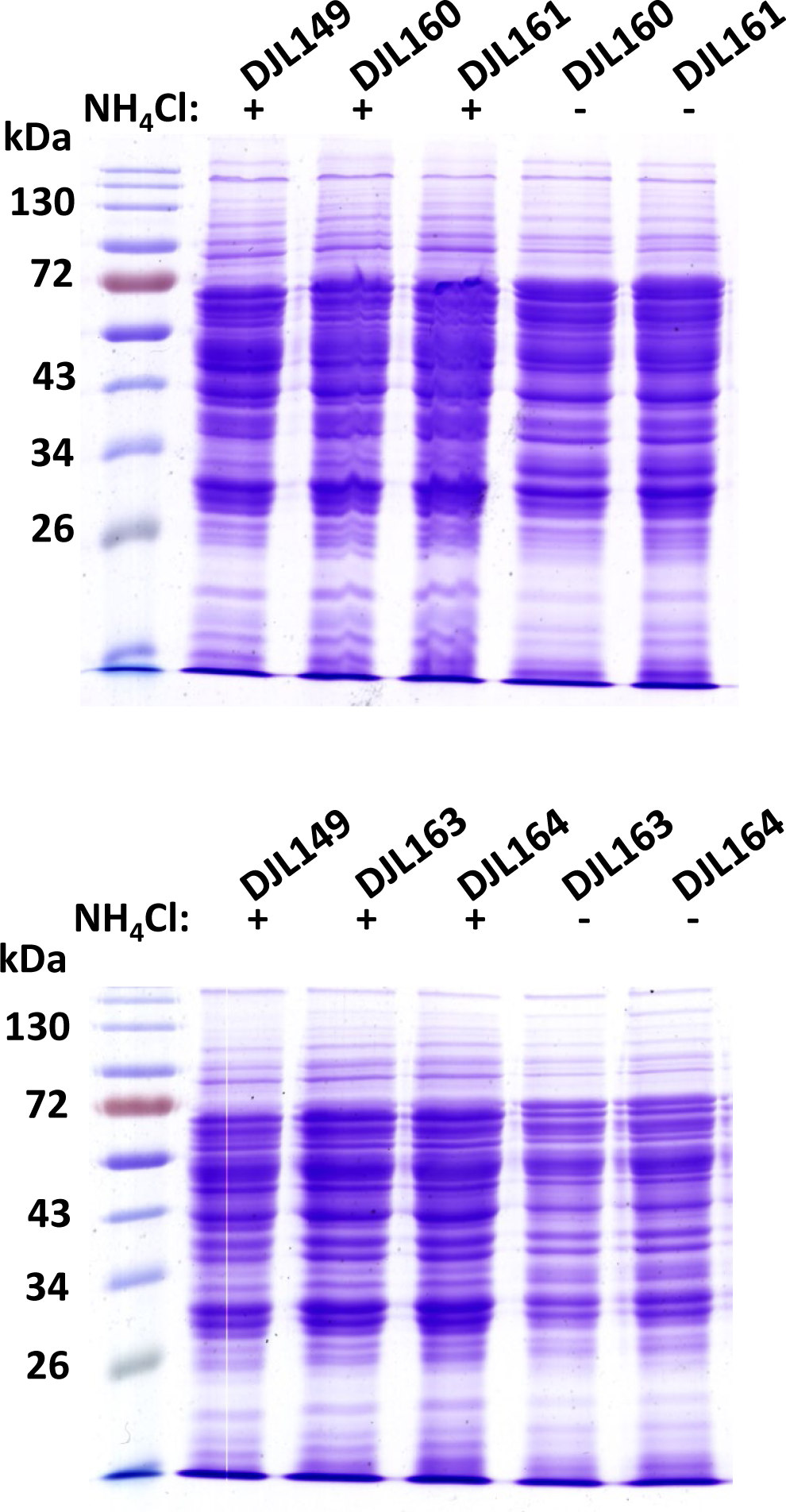
**P**rotein loading control SDS-PAGE of lysates for the Western blots shown in Fig 6B.

**Figure S11.**
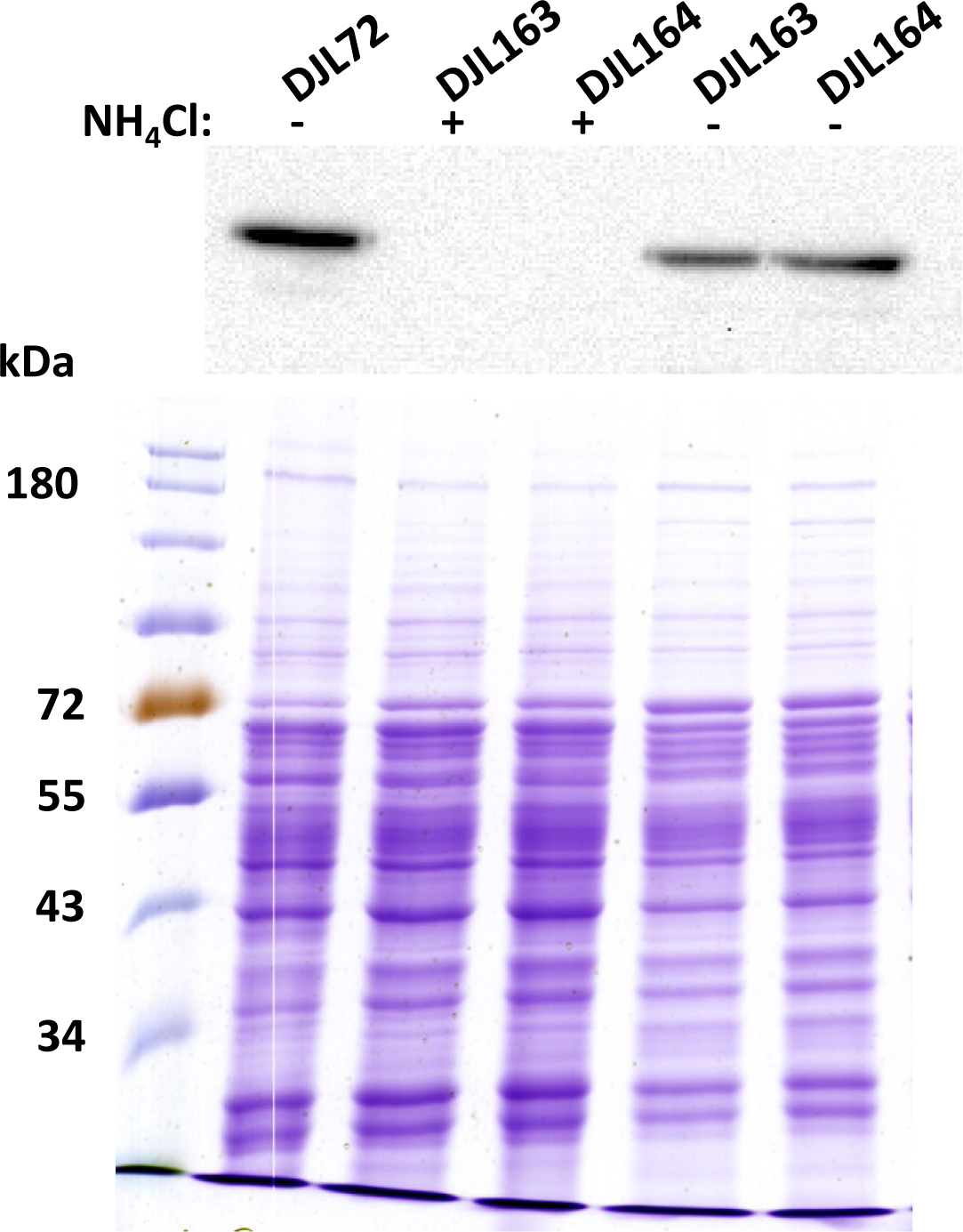
Western blot analysis of NifD abundance in cell lysates of *M. acetivorans* strains DJL163 and DJL164. Strains were grown in HS_DTT_ medium containing 125 mM methanol and 1 mM Na_2_S with or without NH_4_Cl. Protein loading control SDS-PAGE of lysates used for is shown below Western blot.

